# Transcriptionally distinct atypical B cell populations contribute to insulin autoreactivity and show differential activation, isotype switching, and somatic hypermutation in early-stage type 1 diabetes individuals

**DOI:** 10.64898/2026.09.30.754202

**Authors:** Lindsay E. Bass, Landon M. Clark, Bryan Han, Daniel J. Moore, Rachel H. Bonami

**Author notes:** Correspondence: Rachel H. Bonami, Department of Medicine, Vanderbilt Health, Medical Center North T3113, 1161 21st Avenue South, Nashville, TN 37232.

## Abstract

Autoreactive B cells must escape immune tolerance mechanisms to drive autoimmune disease, potentially through alternative paths to memory and antibody-secreting cell formation. “Atypical” B cells represent one such alternative pathway and are expanded in several human autoimmune diseases. Atypical B cells display phenotypic heterogeneity, but how this connects with differences in transcriptional programming, isotype switching and somatic hypermutation is incompletely understood in type 1 diabetes (T1D). Screening efforts to detect T1D prior to clinical diabetes onset affords an opportunity to study B cells early in immunopathogenesis, in which they primarily function as antigen-presenting cells. Prior evidence points to atypical B cell populations as reservoirs for islet autoreactivity in human T1D. We therefore developed sorting strategies to enrich for atypical and other autoreactive-prone B cell populations from the peripheral blood of early-stage T1D individuals and performed single-cell RNA/BCR-sequencing. We identified four transcriptionally defined atypical B cell clusters, each showing differential upregulation of atypical B cell defining genes. Additional gene expression changes were noted in broadly transcriptionally similar, but phenotypically distinct atypical B cell populations, with differences noted in the average extent of isotype switching and somatic hypermutation. Some atypical B cell populations showed extensive clonal sharing with classical memory or plasmablast subsets, consistent with relationships amongst these subsets. B cells reactive against a key islet autoantigen, insulin, were enriched in select atypical B cell populations. These findings highlight specific atypical B cell populations and features with potential value as biomarkers of disease progression or response to immunotherapy in T1D, and potentially other autoimmune diseases.

## 1 Introduction

While several immunotherapy agents show promise in limiting immune-mediated beta cell destruction in type 1 diabetes (T1D), no FDA-approved therapy provides sustained protection against clinical diabetes onset (1). Immunotherapy clinical trials have often shown heterogenous responses with only a subset of individuals experiencing positive outcomes (reviewed in (1)). A deeper understanding of heterogenous immune populations in human T1D - the focus of this study - could be leveraged to inform stratification of individuals for appropriate treatment regimens in the future. For example, with immunotherapies designed to modulate B-T interactions, polyclonal B and T cells reactive against irrelevant antigens obscure our ability to measure what matters - changes in antigen-specific responses. Minimally supervised analysis approaches such as T-REX have successfully extracted vaccine responsive B and T cell populations from the “noise” of total PBMCs using 40 phenotypic markers (2). Given that autoreactive B cells often defy dogmas established in protective immune responses, we developed an integrated strategy combining phenotypic, immune repertoire, and transcriptomic profiling with screening against the key T1D autoantigen, insulin. Our goals were to identify population(s) in which anti-insulin B cells are enriched and to determine which atypical and CD21lo B cell populations associate most closely with features providing evidence of recruitment into immune responses (e.g., clonally expanded, isotype switched, mutated BCRs). More broadly, defining these antigen-specific and response-associated populations may establish additional biomarkers and programs for monitoring disease-relevant immune changes.

Immunotherapies that deplete B lymphocytes (B cells) temporarily preserved beta cell function in T1D clinical trials (3, 4), implicating B cells in human T1D pathogenesis. B cells can differentiate to produce islet autoantibodies used to predict T1D disease risk (5, 6). Studies in the non-obese diabetic (NOD) mouse model of T1D suggest that autoreactive B cells chiefly promote disease via presentation of islet autoantigens to autoreactive T cells (7–12). Germinal center, but not other B cells reactive against the islet autoantigen, insulin, were competent to process and present insulin peptides to a pathogenic T cell clone, providing an example of how B cell programming differentially controls their pathogenic potential (13). T cell interactions also shape islet-reactive B cell programming in T1D, as T cell loss of BCL6 limits anti-insulin B cell differentiation into germinal center and atypical B cell fates (14). This is associated with diabetes protection in an NOD model (14), emphasizing a need to better understand atypical B cells in human T1D.

Atypical B cells are a broad class of distinct B cell subsets for which nomenclature can vary in literature. Here, we define the broader term, “atypical B cells” as encompassing atypical memory B cells, age-associated B cells, CD21lo B cells, and double negative B cells (classified into DN1, DN2, and DN3), in line with surface marker and gene expression programs (reviewed in (15, 16)). Atypical B cell features include lack of CD27 expression and/or low CD21 expression, in addition to expression of genes including ITGAX, TBX21, ZEB2, and FCRL5 (15). While atypical B cells are identified in healthy individuals, expansion of specific atypical B cell populations has been shown in systemic lupus erythematous (SLE), rheumatoid arthritis (RA), systemic sclerosis, multiple sclerosis, neuromyelitis, and autoimmune liver disease (15, 17–22). Atypical B cells contribute to autoimmune pathogenesis in the MRL/lpr murine model of lupus (23).

Studies from our lab and others show atypical B cell subsets harbor insulin autoreactive B cells in human T1D (24–27). Insulin is a critical autoantigen in both human and murine T1D (7, 28–30). To circumvent the technical challenges related to studying rare, antigen-specific B cells in humans, we have also studied anti-insulin B cells in T1D-prone NOD mice using B cell receptor (BCR) transgenes to enrich this population. Specifically, VH125^SD^.NOD mice have an increased frequency (1-3%) of anti-insulin B cells and develop accelerated diabetes onset relative to WT.NOD mice (31). As an additional benefit, the VH125^SD^.NOD model enables anti-insulin B cell study in the complex microenvironment of the pancreas, which cannot be practically biopsied in humans. Whereas few anti-insulin B cells undergo proliferation in the spleen, nearly 20% of anti-insulin B cells have recently proliferated in the pancreas of VH125^SD^.NOD mice (14), consistent with their pathogenic role as antigen presenting cells in T1D (7, 12). Anti-insulin B cells in VH125^SD^.NOD mice skew toward the CD11c+ CD11b+ and CD11c+ T-bet+ atypical B cell subsets in the pancreas, relative to non-insulin-binding B cells, and compared to splenic B cell populations (14). Given the pathogenic potential of atypical B cells and challenges classifying these cells in the field (15, 16), we designed a strategy to enrich for phenotypic atypical B cells to identify previously unrecognized atypical B cell transcriptional heterogeneity in human T1D.

## 2 Materials and methods

### 2.1 Participant selection and clinical information

All participants were recruited at Vanderbilt University Medical Center via the TrialNet Pathway to Prevention study and met inclusion/exclusion criteria listed in **Supplemental Table S1**. Male and female individuals were represented in this study (63% female); participant demographics are shown in **Supplemental Table S2**. Non-identifying participant metadata were managed using REDCap software system (Vanderbilt) (32).

### 2.2 Study Approval

These studies were approved by the University of Miami and Vanderbilt Institutional Review Boards. Written informed consent was received prior to participation. Ancillary Study Approval was granted by Type 1 Diabetes TrialNet.

### 2.3 Sample collection and processing

Peripheral blood mononuclear cells (PBMCs) were obtained by collecting whole blood via peripheral venipuncture into Vacutainer CPT Mononuclear Cell Preparation Tubes with sodium citrate (BD). Cells were washed in phosphate buffered saline (PBS) and red blood cells were lysed using ACK Lysis Buffer (Gibco). Cells were washed twice more in PBS and counted. Cells were resuspended and cryopreserved in 10% dimethyl sulfoxide (DMSO) in heat-inactivated fetal bovine serum (FBS) at a rate of -1°C/minute until they reached a temperature of -80°C for 24-72 hours, after which they were transferred to liquid nitrogen storage.

### 2.4 Flow cytometry staining and sorting

PBMCs were rapidly thawed, washed in 1X PBS, and resuspended in flow cytometry staining buffer (1X PBS, 0.02% EDTA, 1% heat-inactivated FBS). Cells were stained on ice with antibodies CD19-AF488 (1:200, HIB19, BioLegend #302219), CD27-PE Dazzle (1:200, O323, BioLegend #302844), CD21-PCPCy5.5 (1:100, Bu32, BioLegend #354908), CD24-APCCy7 (1:100, ML5, BioLegend #311131), CD38-APC (HIT2, BioLegend #303510), and viability dye BV510 (1:1000, Tonbo). Each sample was also stained with a Sample Tag antibody from the BD Single-Cell Multiplexing Kit (Sample Tags #1-8) (BD #633781) to support demultiplexing of the four flow cytometry sort populations that were combined for Rhapsody HT Express Single-Cell profiling. CD19+ viable cells from each participant were flow cytometry sorted into four subsets, named based on sorted populations identified in the representative gating scheme shown in **Supplemental Figure S1**, as follows: *Sort Pop P5*: CD27+ CD21lo, *Sort Pop P6*: CD27+ CD21+, *Sort Pop P8:* CD27-CD21lo CD24lo CD38-, and *Sort Pop P9:* CD27-CD21lo CD24+ CD38+/ CD24+ CD38-/ CD24-CD38+, using a BD FACSAria III sorter in the VUMC Flow Cytometry Shared Resource.

### 2.5 Analysis of differential gene expression (RNA-seq) in single-cell data

To ensure that approximately the same number of cells were included for each BD Rhapsody lane, four B cell populations, Sort Pop 5, Sort Pop 6, Sort Pop 8, and Sort Pop 9 as defined above, were combined. Each of the four Sort Pops for a given lane were isolated from four individual participants stained with separate multiplexing antibodies to support demultiplexing. Combined samples, as outlined in **Supplemental Table S3**, were delivered to the VANTAGE Shared Resource for library preparation and sequencing. The BD Rhapsody HT Express Single-Cell Analysis System was used to create whole transcriptome, human BCR, and Sample Tag libraries using the Ab-O Single Cell Sample Multiplexing Kit, BD Rhapsody cDNA Kit, BD Rhapsody Enhanced Cartridge Reagent Kit V3, BD Rhapsody WTA Amplification Kit, BD Rhapsody TCR/BCR Next Amplification Kit, and BD Rhapsody Dual Index Kit (BD Genomics) according to the manufacturer’s instructions. A target of 50,000 reads per cell was sequenced using an Illumina NovaSeq X Plus (25B) PE150. Data were de-multiplexed (based on hashtag antibody staining) and processed via the Seven Bridges Genomics Platform using the BD Rhapsody Sequencing Analysis Pipeline (v2.2.1 and v2.3) using GRCh38 as a reference.

Custom R scripts were developed to enable simultaneous single-cell transcriptomic (RNA-seq) and immune repertoire (BCR-seq) data analysis using the Seurat package (v5.1.0), the Immcantation/Change-O suite (33), and IMGT-HighV-QUEST (34).

Cells with fewer than 200 RNA features and/or that contained >30% mitochondrial genes were removed. This 30% threshold was selected based on the bimodal distribution of mitochondrial gene percentages observed within each BD Rhapsody lane, with the vast majority of cells falling below this threshold (data not shown). RNA-seq data were normalized and scaled using the SCTransform function. Data from individual lanes were integrated using SelectIntegrationFeatures, PrepSCTIntegration, FindIntegrationAchors, and IntegrateData functions in Seurat. Immunoglobulin VH, Vκ, Vλ genes were regressed out to prevent them from driving cluster assignments using FindNeighbors and FindClusters. B cell subset identities were manually assigned to individual Seurat-determined clusters based on transcriptional profiles that were consistent with other studies defining these populations (27, 35–38), and with the phenotypic identity of the sorted populations from which each cell came.

For all differentially expressed gene (DEG) analyses, VH, Vκ, Vλ, JH, Jκ, Jλ, Cκ, Cλ, genes were omitted as uninteresting sources of biological variation, given our separate analysis of immune repertoire features. To differentiate each B cell subset/cluster/phenotype from all other cells in the dataset being evaluated, the Seurat FindAllMarkers function was used to identify DEGs (fold change > 1.5, padj < 0.05, and expressed in > 30% of the cells in either group, using the default non-parametric Wilcoxon rank-sum test). For comparisons across sort phenotypes within a given transcriptional cluster, groups with n < 10 cells were excluded from analysis. Venn diagrams showing shared DEGs were generated using the VennDetail package in R.

Gene set enrichment analysis (GSEA) was performed on ranked DEG lists that met the following criteria: padj < 0.05, fold change > 1.5, and expressed by >30% of the cells. HALLMARK gene sets were retrieved from MSigDB using R package msigdbr(species = “Homo sapiens”, category = “H”). GO BP B cell pathway gene sets were retrieved from MSigDB using R package msigdbr(species = “Homo sapiens”, category = “C5”, subcollection = “GO:BP”). The following terms were queried: “B_CELL”, “ANTIBODY”, “IMMUNOGLOBULIN”, “PLASMA_CELL”, “PLASMABLAST”, “GERMINAL_CENTER”, “GERMINAL”, “SOMATIC_HYPERMUTATION”, “AFFINITY_MATURATION”, “CLASS_SWITCH”, “BCR”, “B_LYMPHOCYTE”, “B_CELL_RECEPTOR”, “B_CELL_ACTIVATION”, “B_CELL_DIFFERENTIATION”, “B_CELL_PROLIFERATION”, “MEMORY_B_CELL”, “ANTIGEN_PROCESSING”, which returned the list of gene sets in **Supplemental Table S5.** GSEA was performed on the ranked gene lists using the R package fgsea, which returns normalized enrichment score (NES), false discovery rate (FDR) q values, list size, and leading-edge genes for each pathway. Gene ratio was calculated as the number of leading-edge genes/ size of the pathway.

### 2.6 Immune repertoire analysis of BCR-seq single-cell data

For VDJ analysis, the BD Rhapsody Sequencing Analysis Pipeline was used to filter on cells with high confidence VDJ calls and remove BCRs that were unproductive, not in-frame, or lacked CDR3s and assign V, D, and J gene segment, and isotype. Clonotype assignments were initially made using DefineClones.py from changeo (which defaults to immunoglobulin heavy chain analysis only), and were further refined to require the same 1) Vκ/Vλ gene and 2) Igκ/Igλ junction length. We defined clonally expanded B cells as n ≥ 2 clones, given that ∼ 10,000 cells were profiled per participant with most BCRs representing singleton clonotypes. VDJ data were run through IMGT HighV-QUEST (Alamyar et al 2012) which was used to determine % somatic hypermutation in VH and VL gene segments.

VH and VL gene usage was compared across atypical and memory B cell subsets at the clonotype level. Cells were filtered to those with paired heavy and light chain V gene calls. Clonotypes observed in multiple subsets were counted independently. Only V genes that appeared in n > 5 clonotypes were included in the analysis. A 2×2 contingency table was constructed comparing clonotype counts using each V gene versus all other V genes, within the subset of interest versus all other subsets pooled. Enrichment or depletion was assessed using Fisher’s exact test with Benjamini-Hochberg correction. Log2 fold-change was calculated as the ratio of V gene usage within the subset versus all other subsets, with a pseudocount of 0.25% added to both terms to stabilize estimates near zero.

Clonotype diversity metrics were calculated using participant-specific clonotype abundances using the Shannon diversity index and the Gini-Simpson index in the diversity function from the R package vegan. Pielou’s evenness was calculated by dividing the Shannon diversity index by richness, or the log(number of observed clonotypes). Only participants that had n ≥ 20 cells in all groups were included in the analyses.

Clonotype sharing was analyzed within each participant. Subsets containing n < 20 cells were excluded, and only expanded clones (n ≥ 2) were evaluated. The presence of clonotypes within each subset was compared on an individual participant level across all subsets and the number of shared clones was determined.

### 2.7 Recombinant monoclonal antibody (mAb) expression

V(D)J and VJ nucleotide sequences from the BCR-seq data were expressed as recombinant mAb by *in silico* translating V(D)J and VJ nucleotide sequences into amino acid sequences and were designed with human IgG1 (heavy chain) or Igκ or Igλ (light chain) constant domains. Recombinantly expressed mAbs were protein A purified and dialyzed into 1X PBS by GenScript.

### 2.8 Enzyme-linked immunosorbent assay (ELISA)

MaxiSorp 384-well ELISA plates were coated with 1 μg/ml human insulin (Sigma #I2643) in borate buffered saline (pH ∼8.2) overnight at 37°C and washed 5X with 1/2X PBS. Plates were blocked with 2% bovine serum albumin in 1X PBS for 1-2 hours at room temperature (RT), then washed 5X with 1/2X PBS. Experimental and control mAbs were diluted to 1 μg/ml in PBS-Tween (PBS-T), transferred to the ELISA plate and incubated 1 hour at RT. Plates were washed 5X with 1/2X PBS. Bound mAb was detected by goat anti-human IgG-HRP (SouthernBiotech #2040–05) diluted 1:2500 in PBS-T, incubated 30 min at RT. Plates were washed 5X with 1/2X PBS, TMB Ultra substrate (Thermo Fisher #34029) was added, and plates were read at O.D. 370 nm using a BioTek Synergy LX multi-mode plate reader with a max signal detection limit of O.D. = 3.

### 2.9 Statistical analysis

Standard statistical tests utilized for each experiment are indicated in the corresponding figure legends and significance values were calculated using stats package in base R v4.5.3 or GraphPad Prism v10.4.1 (GraphPad Software).

## 3 Results

### 3.1 CD27+ CD21lo atypical B cells are distributed among four transcriptionally defined atypical B cell clusters, while CD27-CD21lo CD24-CD38-DN2 B cells are predominant in a single cluster

Transcriptional heterogeneity is observed in atypical B cells in chronic infection and other autoimmune settings (37). To enrich for rare atypical B cell populations in the peripheral blood of early-stage T1D individuals, we flow cytometry sorted CD21lo atypical memory and plasmablasts (CD27+ CD21lo, “Sort Pop P5”), classical memory (CD27+ CD21+, “Sort Pop P6”), double negative 2-like (CD27-CD21lo CD24lo CD38-, “Sort Pop P8”), and CD21lo naïve/transitional (CD27-CD21lo CD24+ CD38+/CD24+ CD38-/ CD24-CD38+, “Sort Pop P9”) B cell subsets, as shown in **Figure 1A**, with representative flow cytometry sort plots shown in **Supplemental Figure S1**. We then performed single cell RNA-seq and BCR-seq, as in Methods, to evaluate transcriptional and BCR repertoire differences among these different B cell subsets in human T1D. Data were integrated and Seurat identified 18 distinct transcriptional clusters (agnostic to phenotypic sort population metadata) (**Figure 1B**), with cells from several of the four phenotypically defined sort populations falling into multiple regions on the UMAP, as expected, given the anticipated heterogeneity within each population (**Figure 1C**). We defined Seurat clusters as specific immune cell subsets based on expression of key genes (**Figure 1D**) and the top 10 upregulated genes per cluster (**Supplemental Figure S2**). Individual participants (**Supplemental Figure S3A**) and different BD Rhapsody sequencing lanes (**Supplemental Figure S3B**) were distributed across transcriptionally defined clusters.

**Figure 1.**
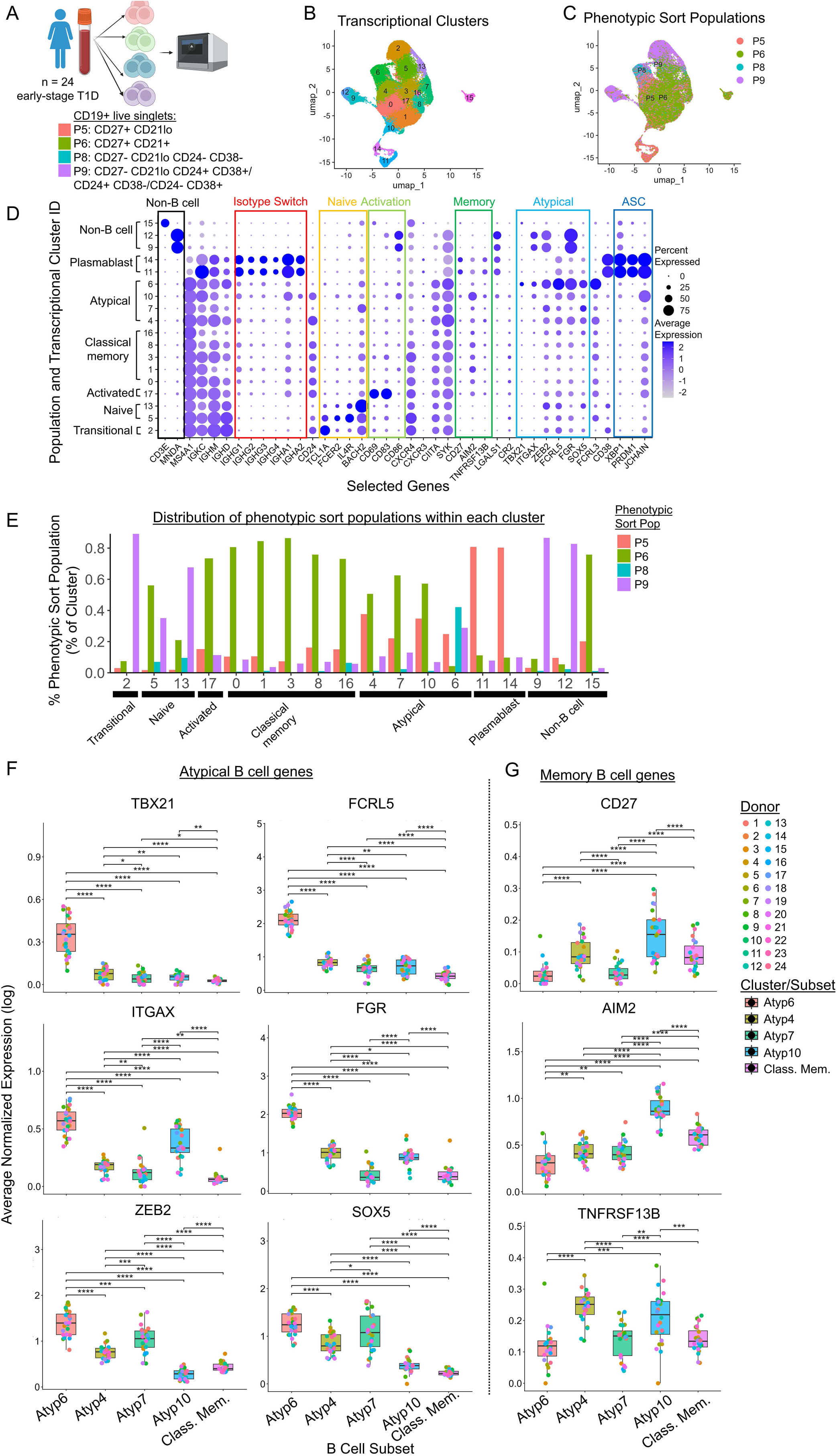
Atypical B cell subsets vary in phenotype and differentially express atypical and memory B cell genes. **A)** Phenotypic B cell subsets were flow cytometry sorted from n = 24 early-stage T1D individuals’ PBMCs, amongst CD19+ live singlets, as follows: (P5) CD27+ CD21lo, (P6) CD27+ CD21+, (P8) CD27-CD21lo CD24-CD38-, and (P9) CD27-CD21lo CD24+ CD38+/CD24+ CD38-/CD24-CD38+ (see representative gating in **Supplemental Figure S1**). Sorted populations (stained with separate hash antibodies) were then combined as in **Supplemental Table S3** and profiled by single-cell RNA-seq/BCR-seq (and CITE-seq for sample multiplexing) with quality control filtering and hash demultiplexing as in Methods. **B)** Seurat identified n = 18 transcriptionally distinct clusters within the integrated dataset of n = 77,031 total cells. **C)** UMAP depicts phenotypically sorted B cell populations. **D)** Transcriptionally defined clusters were broadly assigned as transitional, naïve, activated, classical memory, atypical, plasmablast, or non-B cell based on expression of the indicated selected genes shown, categorized as indicated by the colored rectangles. **E)** The distribution of phenotypic sort populations within each transcriptional cluster was calculated. The Non-B cell subset was excluded from all subsequent analyses (n = 71,566 total B cells remained). **F-G)** Average normalized expression (log) of **F)** atypical B cell genes, TBX21, ITGAX, ZEB2, FCRL5, FGR, SOX5 and **G)** memory B cell genes, CD27, AIM2, and TNFRSF13B are shown for each participant across individual atypical (Atyp) B cell clusters [Atyp6 (n = 4,481 cells), Atyp4 (n = 6,968 cells), Atyp7 (n = 4,252 cell), Atyp10 (n = 2,199 cells)], and combined classical memory clusters (Class. Mem., a combination of Clusters 0, 1, 3, 8, and 16 shown in Figure 1B**/D**, n = 32,311 cells). Freidman test with paired Wilcoxon signed-rank test (Benjamin Hochberg correction), **** p < 0.0001, *** p < 0.001, ** p < 0.01, * p < 0.05, all other comparisons are not significant.

The transcriptionally defined B cell subsets largely aligned with the expected phenotypic sort population identities, with dominant phenotypically defined populations observed in many of the clusters (e.g., transitional, classical memory, and plasmablast clusters) (**Figure 1E**). Individual participant-level distributions for transcriptionally defined clusters (**Supplemental Figure S4A**) and phenotypic sort populations (**Supplemental Figure S4B**) identified representation of these clusters and phenotypes within each individual, with some variation across individuals. Heterogeneity in phenotypically defined populations was noted within specific transcriptionally defined atypical B cell clusters (**Figure 1E**). Atypical Cluster 6 was the only cluster in which an abundant population of phenotypic “DN2-like” B cells were observed, which also contained other phenotypically defined populations at >20% of total cells within this cluster (**Figure 1E**). Of note, our gating strategy to capture DN2 B cells was defined as CD19+ CD27-CD21lo CD24-CD38-, which did not require IgD negativity, as in the classic DN2 B cell gating scheme (CD27-IgD-CD21lo/-CD24-CD38-) (39), thus we will refer to this phenotypic population as “DN2-like”. Overall, these data reveal the transcriptional and phenotypic complexity of atypical B cells in T1D.

### 3.2 Transcriptionally defined atypical B cell clusters show differences in expression of several classical atypical and memory B cell genes

Clusters 4, 7, 10, and 6 were defined as atypical B cell populations based on expression of the following genes: TBX21 (T-bet), ITGAX (CD11c), ZEB2, FCRL5, FGR, and SOX5. Observable differences in expression of classical atypical B cell genes across atypical Clusters 4, 7, 10 and 6 (**Figure 1D**) led us to further investigate gene expression of atypical and memory B cell genes between the atypical B cell clusters at the participant level, with classical memory B cells (combining Clusters 0, 1, 3, 8, and 16) included as a reference. Cluster 6 B cells expressed the highest levels of atypical B cell genes TBX21 (T-bet), ITGAX (CD11c), ZEB2, FCRL5, FGR, and SOX5 compared to Atypical Clusters 4, 7, 10 and classical memory (**Figure 1F**). ITGAX expression was highest in Clusters 6 and 10, with the highest level of ZEB2 expression observed in Clusters 6 and 7 (**Figure 1F**). Cluster 10 was the only atypical cluster that showed decreased expression of ZEB2 compared to classical memory (**Figure 1F**). Cluster 4 upregulated FCRL5 compared to Clusters 7, 10, and classical memory, although not to the extent of Cluster 6 (**Figure 1F**). FGR and SOX5 were upregulated in all atypical clusters compared to classical memory, except Cluster 7, in which SOX5 was elevated but FGR was not (**Figure 1F**). We also evaluated expression of memory-defining B cell genes included CD27, AIM2, and TNFRSF13B (TACI), which were expressed by the classical memory B cells, as expected (**Figure 1G**). Atypical Cluster 10 expressed increased levels of memory genes CD27, AIM2, and TNFRSF13B compared to classical memory (**Figure 1G**). These data define four distinct atypical B cell populations that differentially upregulate genes classically used to define atypical and memory B cells, suggesting a single transcriptional definition of atypical B cells fails to identify distinct atypical B cell subsets in T1D.

### 3.3 Distinct atypical B cell clusters show transcriptional differences in genes related to B cell signaling and activation

To evaluate whether the atypical B cell subsets identified in human T1D may be transcriptionally wired for distinct B cell functions, we performed differential gene expression analysis to identify differentially expressed genes (DEGs) compared to classical memory B cells as a reference, to see which, if any DEGs were shared or unique across each atypical B cell cluster. Only one gene, FCRL5, was significantly upregulated across all four atypical B cell subsets compared to classical memory (**Figure 2A, Supplemental Table S4**), and only four genes (CXCR4, TXNIP, ZFP36L1, ZFP36L2) were significantly downregulated across all atypical clusters compared to classical memory (**Figure 2B, Supplemental Table S4**). Upregulation of FCRL5 and downregulation of CXCR4 suggest distinct regulation of atypical B cells compared to classical memory B cells.

**Figure 2.**
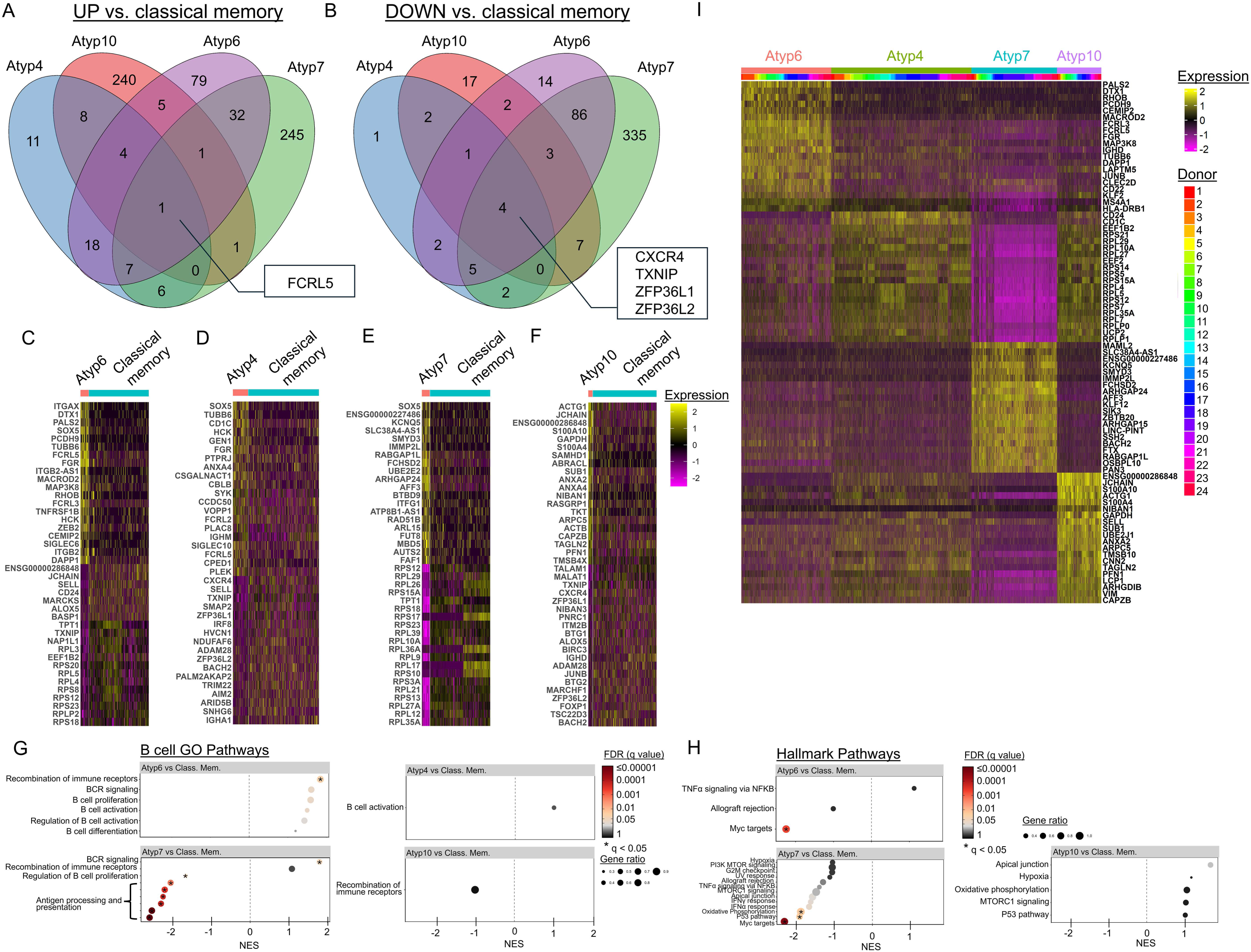
Atypical B cell clusters show distinct gene expression profiles from classical memory and from one another. B cells were isolated and transcriptionally profiled as in Figure 1. DEGs were identified by FindMarkers according to the following criteria: padj < 0.05, fold change > 1.5, and expressed by >30% of the cells in at least one of the comparator groups. Immunoglobulin V and J genes, and light chain constant region genes were removed from the DEG list as an uninteresting source of biological variation. Venn diagrams note the number of shared and unique **A)** upregulated or **B)** downregulated genes among atypical (Atyp) subsets compared to classical memory, with the genes shared across all atypical subsets listed in white boxes. Heatmaps show the top 20 upregulated and top 20 downregulated genes in **C)** Atyp6, **D)** Atyp4, **E)** Atyp7, and **F)** Atyp10 compared to Classical Memory (a combination of Clusters 0, 1, 3, 8 and 16 shown in Figure 1B**/D**). Gene set enrichment analyses (GSEA) were performed on ranked differentially expressed gene lists (Atyp6 vs Class. Mem., Atyp4 vs Class. Mem., Atyp7 vs Class. Mem., Atyp10 vs Class. Mem.) that met the criteria listed above. GSEA was performed on the ranked gene lists against the MSigDB **G)** GO BP B cell pathways (listed in **Supplemental Table S5**) and **H)** HALLMARK pathways, as in Methods. Pathways were filtered on normalized enrichment score (NES) > 1 or NES < -1. Color scale notes false discovery rate (FDR) q values and circle size notes gene ratio. **I)** Heatmap shows the top 20 DEGs in each atypical cluster compared to all other atypical clusters combined, identified by FindAllMarkers, which met the following criteria: padj < 0.05, fold change > 1.5, > 30% Subset and participant data are color coded.

The majority of DEGs identified within each atypical B cell cluster (relative to classical memory) were unique to that cluster, with the exception of Atypical Cluster 4, which showed more overlapping than unique DEGs (**Figure 2A**). Atypical Cluster 4 upregulated n = 11 unique genes, Cluster 6 upregulated n = 79 unique genes, Cluster 10 upregulated n = 240 unique genes, and Cluster 7 upregulated n = 245 unique genes compared to classical memory (**Figure 2A**). The top 20 upregulated and downregulated genes for each atypical B cell cluster compared to classical memory are shown as heatmaps (**Figure 2C-2F**). Gene set enrichment analysis (GSEA) was performed for each atypical B cell cluster compared to classical memory against the HALLMARK gene list and a list of GO BP B cell-related gene pathways (**Supplemental Table S5**) from the Molecular Signatures Database. The GO BP BCR recombination pathway was enriched in atypical Cluster 6 compared to classical memory (NES = 1.86, padj. = 0.028) (**Figure 2G**). The GO BP BCR signaling pathway was upregulated in Cluster 7 (NES = 1.74, padj. = 0.027), while GO BP antigen processing and presentation (NES = -2.25, padj. = 0.003) and BCR recombination (NES = -1.67, padj. = 0.033) were downregulated in Cluster 7 compared to classical memory (**Figure 2G**). Leading genes for the upregulated and downregulated gene pathways (NES > 1 or < -1) are listed in **Supplemental Table S6**.

The HALLMARK Myc targets gene pathway was downregulated in atypical Cluster 6 (NES = -2.25, padj. = 0.0007) and atypical cluster 7 (NES = -2.35, padj. = 0.00008) compared to classical memory (**Figure 2H**). Cluster 7 showed downregulation of other HALLMARK metabolic pathways, oxidative phosphorylation (NES = -1.89, padj. = 0.022) and P53 pathway (NES = -1.94, padj. = 0.022) (**Figure 2H**). Differential expression of these gene pathways may suggest differences in metabolic requirements or response to stress in atypical B cell clusters 6 and 7 compared to classical memory. No upregulated or downregulated gene pathways with NES > 1 and q < 0.05 were identified in Clusters 4 or 10 compared to classical memory (**Figure 2H**). Overall, atypical Cluster 4 showed differential expression of the lowest number of genes compared to classical memory (n=72), while Cluster 7 differential expressed the highest number of genes (n=735) compared to classical memory. Overall, these data highlight distinct transcriptional programming across the atypical B cell clusters.

Identification of unique upregulated genes in each atypical cluster compared to classical memory (**Figure 2A**) led us to examine gene expression differences between the atypical clusters directly. This analysis emphasized transcriptional differences of Clusters 6 and 7 and transcriptional similarities of Clusters 4 and 10 (**Figure 2I**). Notably, Atypical Cluster 4 and 10 showed the highest frequency of phenotypic CD27+ CD21lo B cells within each cluster compared to the other atypical B cell and classical memory B cell clusters (**Figure 1E**). Cluster 6 significantly upregulated B cell signaling genes DAPP1, MAP3K8, and JUNB and regulatory genes FCRL3 and CD22 compared to the other atypical clusters (**Figure 2I**). Atypical Cluster 4 upregulated various ribosomal proteins (RPS21, RPL29, RPL10A, RPL27, RPS14, RPS5, RPS15A, RPL4, RPL5, RPS12, RPS7, RPL35A, RPL7, RPLP0, and RPLP1) compared to the other atypical clusters (**Figure 2I**), which may suggest increased protein synthesis or metabolic demand. Atypical Cluster 7 significantly upregulated actin cytoskeleton remodeling gene FCHSD2, which is recruited to BCR aggregates during BCR internalization (40), and endosomal trafficking genes ARHGAP24, ARHGAP15, and RABGAP1L. Atypical Cluster 10 significantly upregulated the T cell costimulatory molecule, CD86 (**Supplemental Table S7**), actin rearrangement genes (ACT1G, ARPC5, and PFN1) and JCHAIN (**Figure 2I**), which supports secreted IgA/IgM antibody dimer/pentamerization. The transcriptional heterogeneity observed across all four atypical clusters suggests they exhibit differences in programming related to B cell signaling and activation.

### 3.4 Transcription factors associated with atypical B cells vary in expression across distinct atypical B cell clusters, and across some phenotypically defined populations within the same atypical B cell cluster

Dai et al showed that ZEB2 is a critical transcription factor for the differentiation of atypical (“age-associated”) B cells and lupus pathology; single-cell RNA-sequencing of B cells from a patient with new-onset SLE identified upregulated transcription factors in atypical B cells compared to other B cell subsets (41). We observed differential expression of ZEB2 across our atypical B cell clusters (**Figure 1F**); therefore, we examined expression of other atypical B cell-associated transcription factors across the four atypical B cell clusters identified in early-stage T1D individuals, and found different expression patterns (**Figure 3A**), which was not explained by individual participant-level differences (**Figure 3B**).

**Figure 3.**
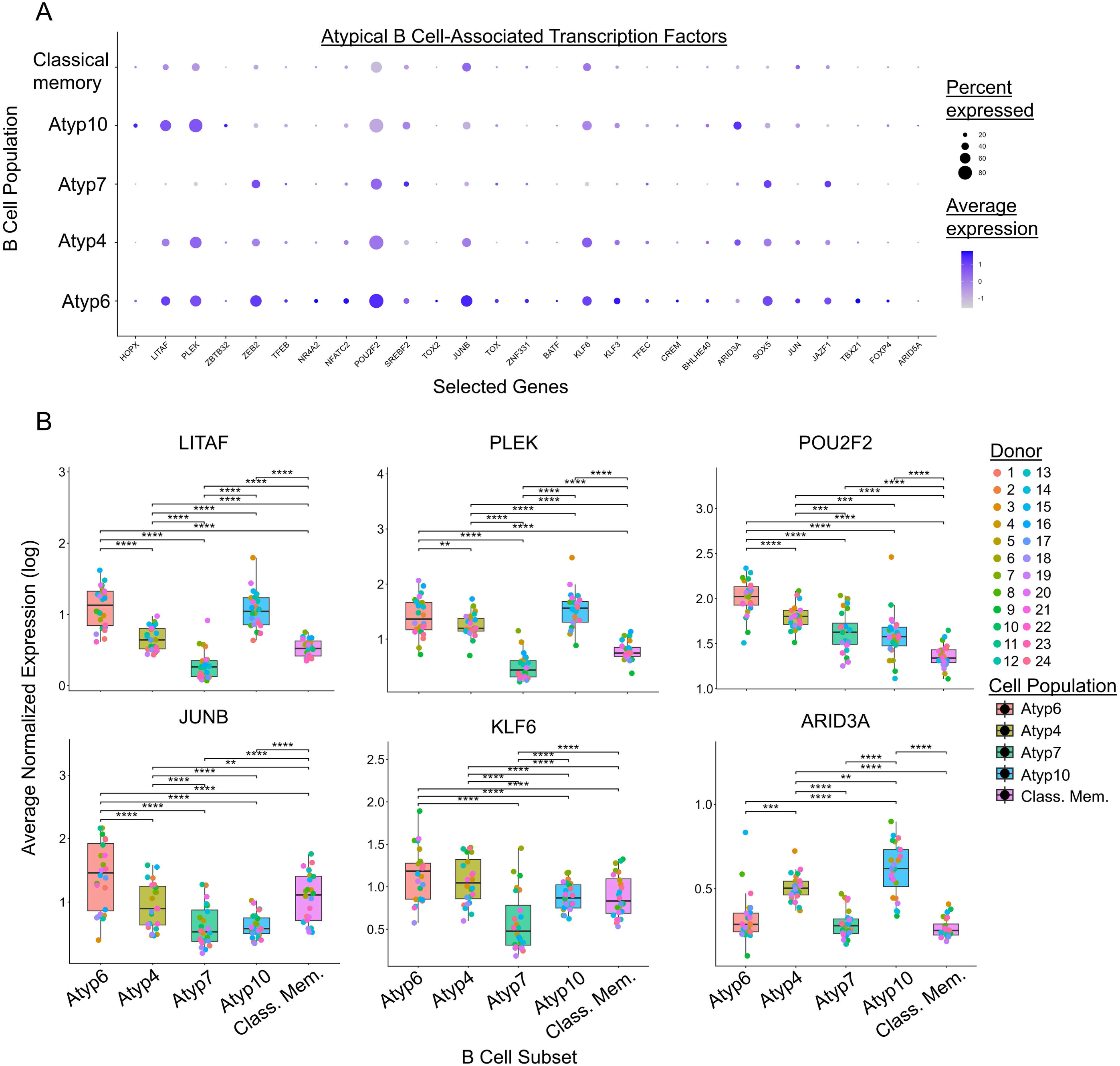
Atypical B cell clusters in early-stage T1D individuals differentially express atypical B cell-associated transcription factor genes. B cells were isolated and transcriptionally profiled as in Figure 1. Atypical B cell clusters and classical memory were evaluated for gene expression of n = 27 transcription factor genes upregulated in atypical B cells compared to other B cell subsets in lupus, with **A)** percent and average expression across the entire population or **B)** average normalized expression (log) of transcription factor genes LITAF, PLEK, POU2F2, JUNB, KLF6, and ARID3A plotted for n = 24 individual participants across atypical clusters (Atyp6, Atyp4, Atyp7, Atyp10), and classical memory (Class. Mem., a combination of Clusters 0, 1, 3, 8 and 16 shown in Figure 1B**/D**). Freidman test with paired Wilcoxon signed-rank test (Benjamin Hochberg correction), **** p < 0.0001, *** p < 0.001, ** p < 0.01, * p < 0.05, all other comparisons are not significant.

Ablation of JUN in cultured human PBMCs impaired double negative CD27^−^IgD^−^CD11c^+^T-bet^+^ B cell formation (41). Cluster 6 showed the highest expression of JUNB, consistent with the enrichment within this cluster of DN2-like phenotypic cells (**Figure 3B**). Constitutive expression of ARID3a in CD19+ B cells in a transgenic mouse model led to production of anti-nuclear autoantibodies and glomerulonephritis, suggesting enhanced expression of ARID3a results in failure to maintain B cell tolerance (42, 43). Cluster 10 showed increased expression of ARID3A (**Figure 3B**), suggesting these cells may be prone to break tolerance and differentiate into antibody secreting cells. Cluster 7 showed decreased expression of LITAF, PLEK, and KLF6 compared to all other atypical clusters (**Figure 3B**), suggesting distinct transcriptional regulation of this cluster compared to the other atypical B cell clusters. Overall, these data highlight differences in transcriptional regulation which could have functional consequences.

Given the phenotypic diversity observed within the atypical B cell clusters (**Figure 1E**), we subset each transcriptional cluster (n = 18) into the four phenotypic sort populations (n = 4), as defined in **Figure 1A** and termed “Cluster_SortPop” here. Cluster_SortPop groups with n < 50 cells were excluded from further analyses, as indicated in **Supplemental Table S8**. We observed subtle differences in the expression of atypical B cell genes across the atypical Cluster_SortPop groups (**Supplemental Figure S5**). Therefore, we evaluated the expression of the atypical B cell-associated transcription factors in **Figure 3** across Cluster_SortPop groups (**Figure 4A**). Cluster 6 showed the most diverse distribution of phenotypic populations within the cluster (**Figure 1E**); therefore, we evaluated participant level expression of transcription factors LITAF, PLEK, ZEB2, POU2F2, JUNB, KLF6, ARID3A, and SOX5. Expression of ZEB2, POU2F2, and JUNB did not significantly differ across sorted populations within Cluster 6 (**Figure 4B**). Atypical Cluster 6 sort population CD27+ CD21lo (6_P5) significantly upregulated KLF6 and ARID3A, but downregulated SOX5 compared to DN2-like sort population CD27-CD21lo CD24-CD38- (6_P8) in Cluster 6 (**Figure 4B**). Average normalized transcription factor expression for the remaining atypical B cell clusters broken down into phenotypic populations are shown in **Supplemental Figure S6**, with few differences noted across phenotypic sort populations within a given transcriptional cluster.

**Figure 4.**
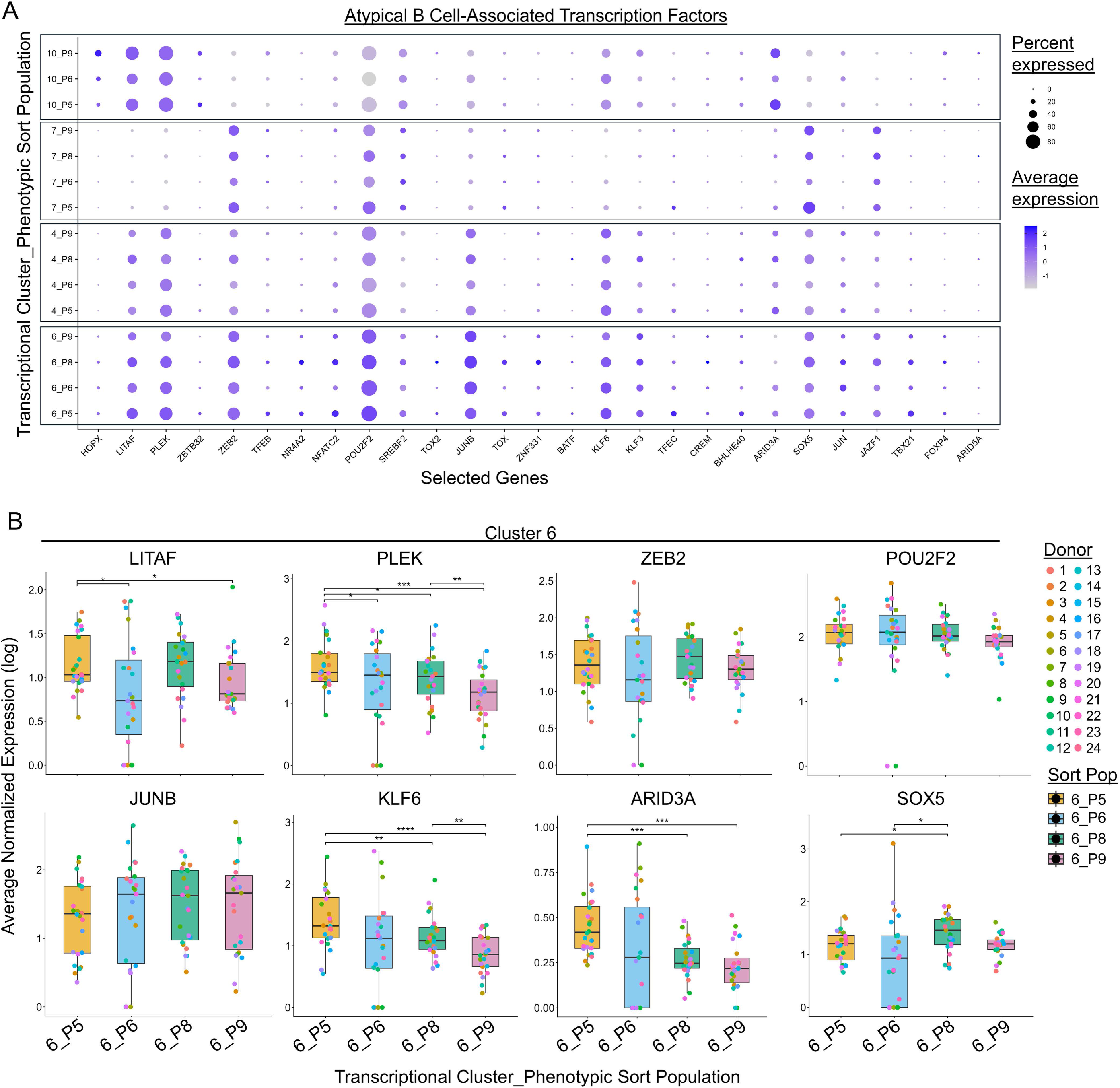
Distinct phenotypic subsets within DN2-like Atypical Cluster 6 differentially express atypical B cell-associated transcription factors, including SOX5 and ARID3A. B cells were isolated and transcriptionally profiled as in Figure 1. Atypical B cell clusters were subclassified by phenotypic sort population. Phenotypic sort populations within transcriptionally defined atypical B cell clusters were evaluated for gene expression of n = 27 transcription factor genes upregulated in atypical B cells in lupus, as in Figure 3**. A)** Percent and average expression across the entire population or **B)** average normalized expression (log) of transcription factor genes LITAF, PLEK, ZEB2, POU2F2, JUNB, KLF6, ARID3A and SOX5 are shown for cluster 6 sort populations (6_P5, 6_P6, 6_P8, 6_P9). In panel B, individual participants (n = 24) are plotted as points. Paired statistical analysis was performed across participants (n = 22) with cells in all four sort populations; two individuals did not have cells in all groups and thus were excluded from statistical analysis. Freidman test with paired Wilcoxon signed-rank test (Benjamin Hochberg correction), **** p < 0.0001, *** p < 0.001, ** p < 0.01, * p < 0.05, all other comparisons are not significant.

### 3.5 Differentially expressed genes are identified among phenotypically distinct B cells in transcriptionally defined atypical B cell clusters

To more thoroughly evaluate heterogeneity within an individual atypical B cell cluster, we performed differential gene expression analysis to identify the top 20 upregulated genes in a given sort population. In Atypical Cluster 4, n = 18 DEGs were identified (**Supplemental Table S9**); only one gene, THEMIS2, was significantly upregulated in Cluster 4 phenotypic population CD27-CD21-CD24-CD38- (4_P8). THEMIS2 has been shown to increase B cell responses to self antigen when presented at low avidity and/or low affinity (44) (**Supplemental Figure S7A**). In Atypical Cluster 6, n = 53 differential upregulated genes were identified (**Supplemental Table S9**). Cluster 6 phenotypic population CD27+ CD21lo (6_P5) upregulated genes including T cell costimulatory molecule CD86 and antigen presentation gene CD1C, while CD27+ CD21+ (6_P6) upregulated CD44, among other genes. DN2-like B cells (6_P8) upregulated genes RHOB, PALS2, RGS2, CEMIP2, and DUSP1 (**Supplemental Figure S7B**). PALS2, also known as MPP6, is an adaptor protein shown to interact with activation induced cytidine deaminase to promote class switching (45). Sort population P9 (6_P9) upregulated CD79B and BACH2, aligning with a more naïve-like population (**Supplemental Figure S7B**). Atypical Cluster 7 showed differential upregulation of n = 18 genes (**Supplemental Table S9**). Cluster 7 sort population CD27+ CD21lo (7_P5) upregulated genes including integrin gene ITGB1, while phenotypic population CD27+ CD21+ (7_P6) upregulated homing and homeostasis gene CXCR4 (**Supplemental Figure S7C**). Atypical Cluster 10 showed differential upregulation of n=32 genes (**Supplemental Table S9**); phenotypic population CD27+ CD21lo (10_P5) upregulated antigen processing and presentation genes, CDC42 and IFI30 (**Supplemental Figure S7D**). IFI30 was significantly upregulated in the CD27+ CD21lo subset compared to other transcriptionally defined memory B cell subsets previously (27). JCHAIN was upregulated in both sort populations 10_P5 and 10_P9 when compared to the remaining atypical B cells (**Supplemental Figure S7D**). These data highlight gene expression differences among distinct phenotypic populations, even within transcriptionally defined atypical B cell clusters.

### 3.6 Atypical CD27+ CD21lo cells found within transcriptionally defined classical memory and activated clusters upregulate atypical genes, including SOX5 and FGR

Atypical CD27+ CD21lo (P5) cells were found among all transcriptionally defined clusters, although at low frequencies within the transitional and naïve clusters, as expected (**Figure 1E**). Therefore, we asked whether the CD27+ CD21lo cells would maintain an atypical B cell transcriptional signature even when found in a transcriptionally “non-atypical” B cell cluster. FGR was upregulated in CD27+ CD21lo (P5) cells relative to other phenotypic populations in Classical Memory Cluster 1 (**Supplemental Figure S8A**) and Classical Memory Cluster 8 (**Supplemental Figure S8B and Supplemental Table S9**). Atypical-defining gene SOX5 was upregulated in CD27+ CD21lo cells relative to other phenotypic populations in Classical Memory Cluster 8 (**Supplemental Figure S8B**), Classical Memory Cluster 16 (**Supplemental Figure S8C**) and Activated Cluster 17 (**Supplemental Figure S9A and Supplemental Table S9**). FCRL5 was significantly upregulated in CD27+ CD21lo cells in Classical Memory Clusters 0, while no atypical genes were upregulated in CD27+ CD21lo cells in Cluster 3 (**Supplemental Figure S8D-E**). CD27+ CD21lo cells were the most prevalent cells among the Plasmablast Clusters 11 and 14 (**Figure 1E**), but this population did not differentially upregulate classical atypical genes across the phenotypic populations (**Supplemental Figure S9E-F**).

Transitional (Cluster 2) and naïve (Clusters 5 and 13) subsets were mainly composed of the phenotypic CD27-CD21lo CD24+ CD38+/ CD24+ CD38-/CD24-CD38+ population (P9) (**Figure 1E**), which did not upregulate classic atypical genes in the transitional cluster but did upregulate SOX5 in Naïve Cluster 13 and FCRL5 in Naïve Cluster 5 (**Supplemental Figure S9B**). These data show that phenotypically defined atypical B cell populations maintain expression of some, but not all, key atypical B cell genes, even when their overall transcriptional profiles match naïve, activated or classical memory populations.

### 3.7 Early-stage T1D individuals show variation in isotype switching across and within transcriptionally defined atypical B cell clusters

Isotype switching reflects B cell activation and is influenced by the cytokine environment. Different effector functions are associated with different isotypes; for example, IgA supports mucosal immunity, while IgG subclasses differ in complement binding and Fc receptor activation: IgG3 > IgG1 > IgG2 (46). We therefore evaluated whether differences in class switching exist across atypical B cell clusters and across different phenotypes within clusters. Transitional and naïve clusters were predominantly non-class switched, in contrast to plasmablasts, which had highest proportion of IgA/IgG-switching, regardless of phenotype, as expected (**Figure 5A**). Classical memory and atypical B cells showed enhanced class switching relative to transitional and naïve clusters, with variation observed across different transcriptionally defined clusters within these broader subset classifications (**Supplemental Figure 10A**). The mean frequency of class switched B cells per participant was significantly decreased in Atypical Clusters 6 (38.7%), Cluster 4 (20.1%), and Cluster 7 (37.0%) compared to Classical Memory (47.6%) (**Supplemental Figure 10B**). Cluster 10 showed a significant increase in class switching compared to classical memory, with a mean 71.5% class switched B cells (**Supplemental Figure 10B**). The frequency of IgG1 B cells was significantly decreased in Atypical Clusters 4 (6.0%) and 7 (16.2%) compared to Classical Memory (21.7%) (**Supplemental Figure S10C**). Atypical Cluster 10 exhibited a significantly higher frequency of switching to IgG2 (11.2%), IgA1 (25.3%), and IgA2 (9.4%) compared to the other atypical clusters and classical memory (**Supplemental Figure S10C**). IgG3 class switching was increased in Cluster 6 (6.5%) compared to the other atypical clusters and classical memory (**Supplemental Figure S10C**).

**Figure 5.**
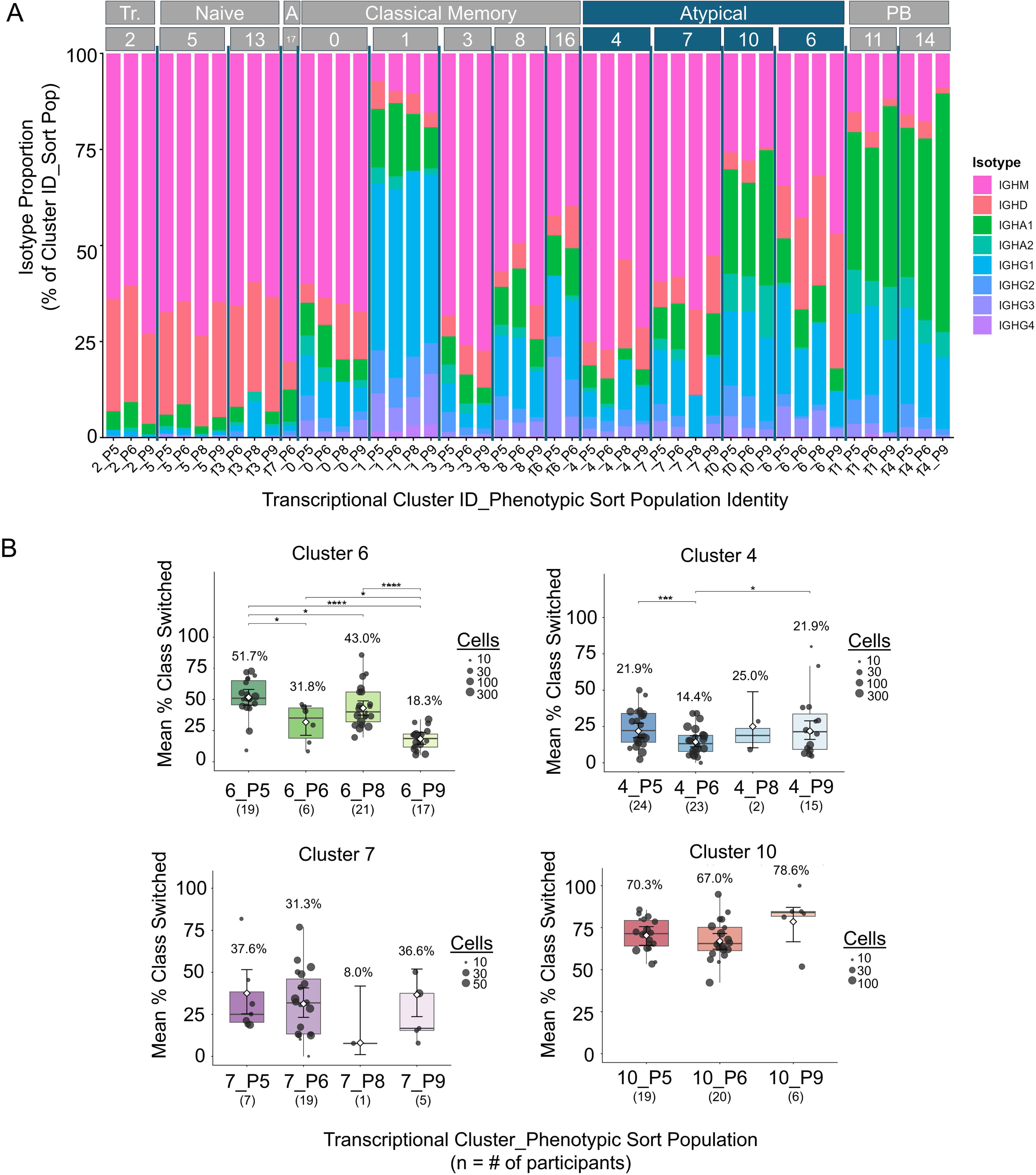
Phenotypic populations within some atypical clusters exhibit different levels of isotype switching. B cells were isolated and transcriptional and immune repertoire profiling was performed as in Figure 1 and Methods. **A)** Isotype distribution was calculated for phenotypic sort populations within transcriptionally defined atypical B cell clusters, with B cell subset and transcriptional cluster IDs shown across the top of the graph. **B)** The average % isotype-switched B cells within each phenotypic sort population is shown for each atypical B cell cluster. Individual participants are plotted as points, with dot size corresponding to number of cells. Phenotypic sort populations with n < 10 cells were excluded from analysis. Boxes show the median and interquartile range of participant means, with whiskers extending to 1.5× IQR. White diamonds and error bars indicate model-estimated marginal means (listed) with 95% confidence intervals. Data were analyzed by a beta-binomial generalized linear mixed model with donor as the random intercept (using the glmmTMB package in R) with Omnibus likelihood-ratio test (LRT) and pairwise comparisons of estimated marginal means (determined by the emmeans package in R and using Wald z-tests) with Benjamini–Hochberg correction, **** p < 0.0001, *** p < 0.001, ** p < 0.01, * p < 0.05, all other comparisons are not significant. Note, for cluster 7, the beta-binomial model did not converge and the fallback analysis (binomial generalized linear mixed model with donor and observation-level random effect) was performed.

Further examination of isotype switching within transcriptional clusters but across phenotypic populations identified substantial heterogeneity across Cluster 6 populations; CD27+ CD21lo SortPop P5 showed 51.7% isotype switching, while CD27-CD21lo SortPop P9 showed 18.3% isotype switching (**Figure 5B**). In contrast to the variation in class switching observed across Cluster 6, isotype switching across phenotypic populations within Atypical Cluster 4 was uniformly low, while Cluster 10 showed uniformly high frequencies of isotype switch across phenotypic populations (**Figure 5B**). Further, B cells with a classical memory phenotype (CD27+ CD21+ SortPop P6) varied widely in the extent of isotype switching across transcriptional clusters (e.g., Cluster 4 14.4% vs. Cluster 10 67.0%, **Figure 5B**). Overall, these data point to potential differences in antigen exposure, T cell help and/or cytokine milieu. This diversity could otherwise be missed when B cell populations are classified solely on the basis of transcriptional or phenotypic definitions. Transcriptional differences may further reveal distinct B cell programming amongst cells expressing the same isotypes.

### 3.8 BCR somatic hypermutation varies across and within atypical B cell clusters within early-stage T1D individuals

Somatic hypermutation is most typically associated with affinity maturation that occurs via T cell selection of clones with favorable mutations in germinal centers; however, mutations can arise through extrafollicular T-dependent and T-independent responses as well (47, 48). Transitional and naïve B cell clusters exhibited little to no somatic hypermutation, with plasmablast clusters showing the highest levels of somatic hypermutation (**Figure 6A and 6B**), as expected based on previous studies of human B cells (49, 50). Atypical Cluster 6 and Cluster 4 showed a significant decrease in mean %VH somatic hypermutation per participant (3.2% and 3.4%, respectively) compared to classical memory (4.8%), with the highest average mutation observed in Atypical Cluster 10 (5.4%) (**Figure 6C**). We also compared mean %VH somatic hypermutation across sorted populations within each atypical cluster. In Atypical Cluster 6, phenotypic CD27+ CD21lo population (6_P5) showed increased %VH somatic hypermutation compared to the other three sort populations (**Figure 6D**). Similarly, phenotypic CD27+ CD21lo population (P5) showed increased %VH somatic hypermutation compared to cluster CD27-CD21lo population (P9) in clusters 4 and 10 (**Figure 6D**). The differences in somatic hypermutation observed here are consistent with differences in B cell-T cell interactions governing mutation and/or selection across atypical B cell populations, which will require functional assays to fully characterize.

**Figure 6.**
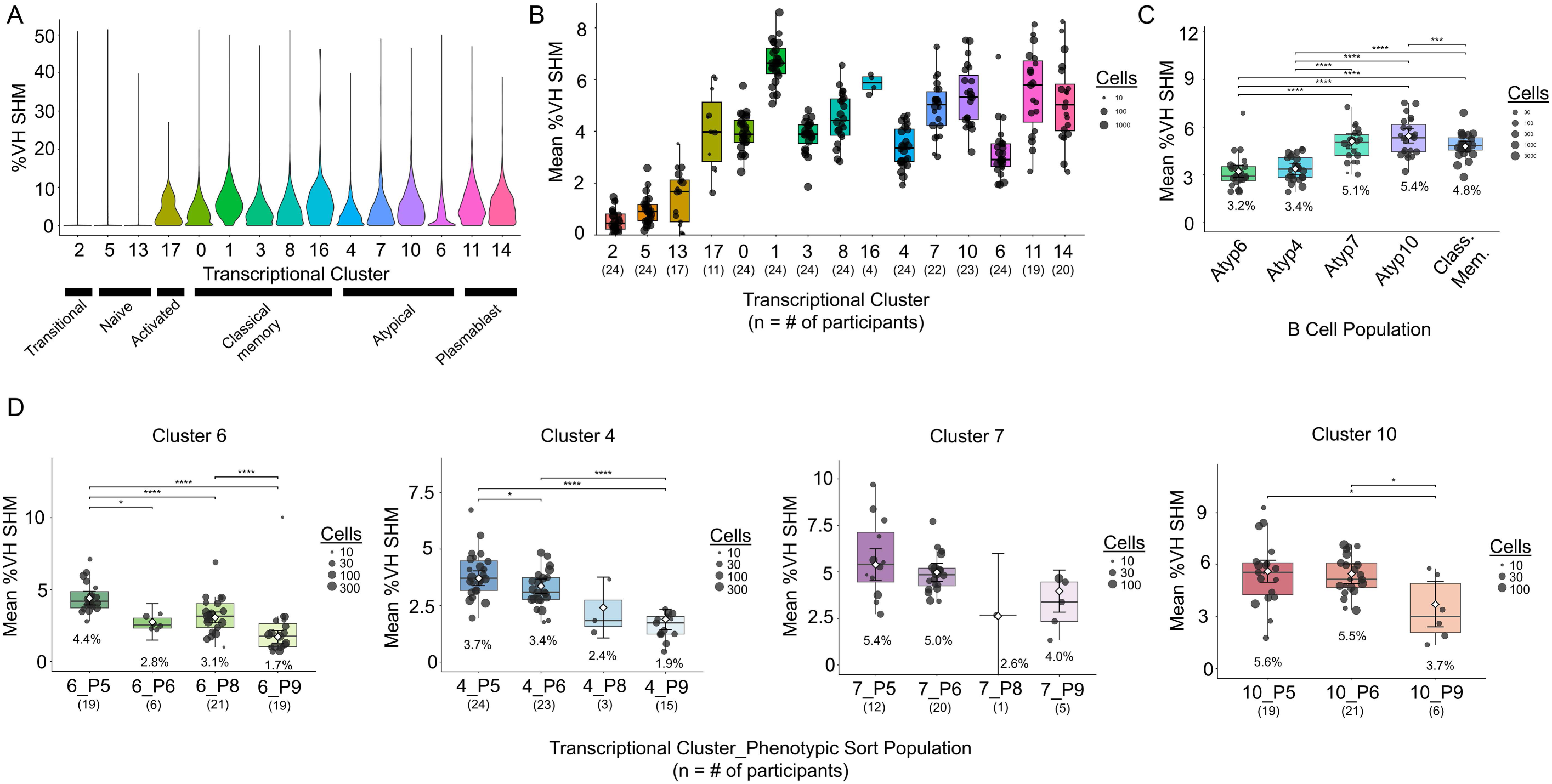
Atypical B cell populations overall, and phenotypic populations within some atypical clusters, exhibit different levels of somatic hypermutation. B cells were isolated and transcriptional and immune repertoire profiling was performed as in Figure 1 and Methods, with IMGT/HighV-QUEST used to determine % VH somatic hypermutation. **A)** % VH somatic hypermutation (SHM) was calculated across all B cell clusters, displayed as violin plots. **B-D)** Individual participants are plotted as points, with dot size corresponding to number of cells. Participant clusters with n < 10 cells were excluded. **B)** Average % VH SHM per participant is shown for each B cell cluster. Boxes show the median and interquartile range of donor means, with whiskers extending to 1.5× IQR. **C-D)** White diamonds and error bars indicate model-estimated marginal means with 95% confidence intervals. Data were analyzed by a precision-weighted linear mixed-effects model (using the Lme4 package in R) on donor-level means (subset as fixed effect, donor random intercept, weights = number of cells), with an omnibus likelihood-ratio test for the effect of subset and pairwise comparisons of estimated marginal means (determined by the emmeans package in R and using Wald t-tests and Satterthwaite degrees of freedom, calculated by the LmerTest package in R) with Benjamini–Hochberg correction. **** p < 0.0001, *** p < 0.001, ** p < 0.01, * p < 0.05; all other comparisons not significant. **C)** Each atypical B cell population was compared against each other, and against classical memory (a combination of Clusters 0, 1, 3, 8 and 16, shown in Figure 1B**/D**). **D)** Each of four phenotypic populations was compared within each transcriptionally defined atypical B cell cluster.

### 3.9 Atypical B cell clusters exhibit increased clonal expansion and decreased diversity compared to classical memory B cells

Given the differences in isotype-switching and somatic hypermutation in different atypical B cell populations observed (**Figures 5-6**), we next examined V gene usage and clonotype diversity. Diverse VH (**Supplemental Figure S11A**) and VL (**Supplemental Figure S11B**) gene usage was observed across atypical and classical memory populations, with differential enrichment of some VH and VL genes noted in some populations (**Supplemental Figure S11C**). Examination of clonotype usage showed that Pielou’s evenness was relatively similar across atypical, classical memory, and plasmablast groups (**Figure 7A**), with values near one suggesting clones within the population are relatively abundant. Some differences were noted in Gini-Simpson diversity (**Figure 7B**), where a Gini-Simpson value of 1 represents maximal diversity. Shannon diversity varied from an index of 3.8 in Atypical Cluster 7 to an index of 7.1 in classical memory, suggesting a high level of diversity among classical memory (**Figure 7C**). Examination of clonal expansion showed limited/no clonal expansion among transitional and naïve B cell subsets, with higher levels of clonal expansion observed in activated, atypical B cells, classical memory, and especially plasmablasts, as expected (**Supplemental Figure S12A**). Specific examination of atypical B cell clusters highlighted increased clonal expansion (defined as in Methods) among Atypical Cluster 4 and Cluster 10 compared to Classical Memory (**Figure 7D**). Of note, clonal expansion only ranged from 0% - 2% across Classical Memory clusters, supporting collapse to one Classical Memory subset (**Supplemental Figure S12A)**. Atypical cluster 10 showed frequency of clonal expansion (7.3%) most similar to the plasmablast subset (12%) (**Figure 7D**). Notably, Cluster 10 located closest to plasmablasts on the UMAP, highlighting transcriptional similarity and a shared proclivity towards enhanced clonal expansion between these populations (**Figure 1B**).

**Figure 7.**
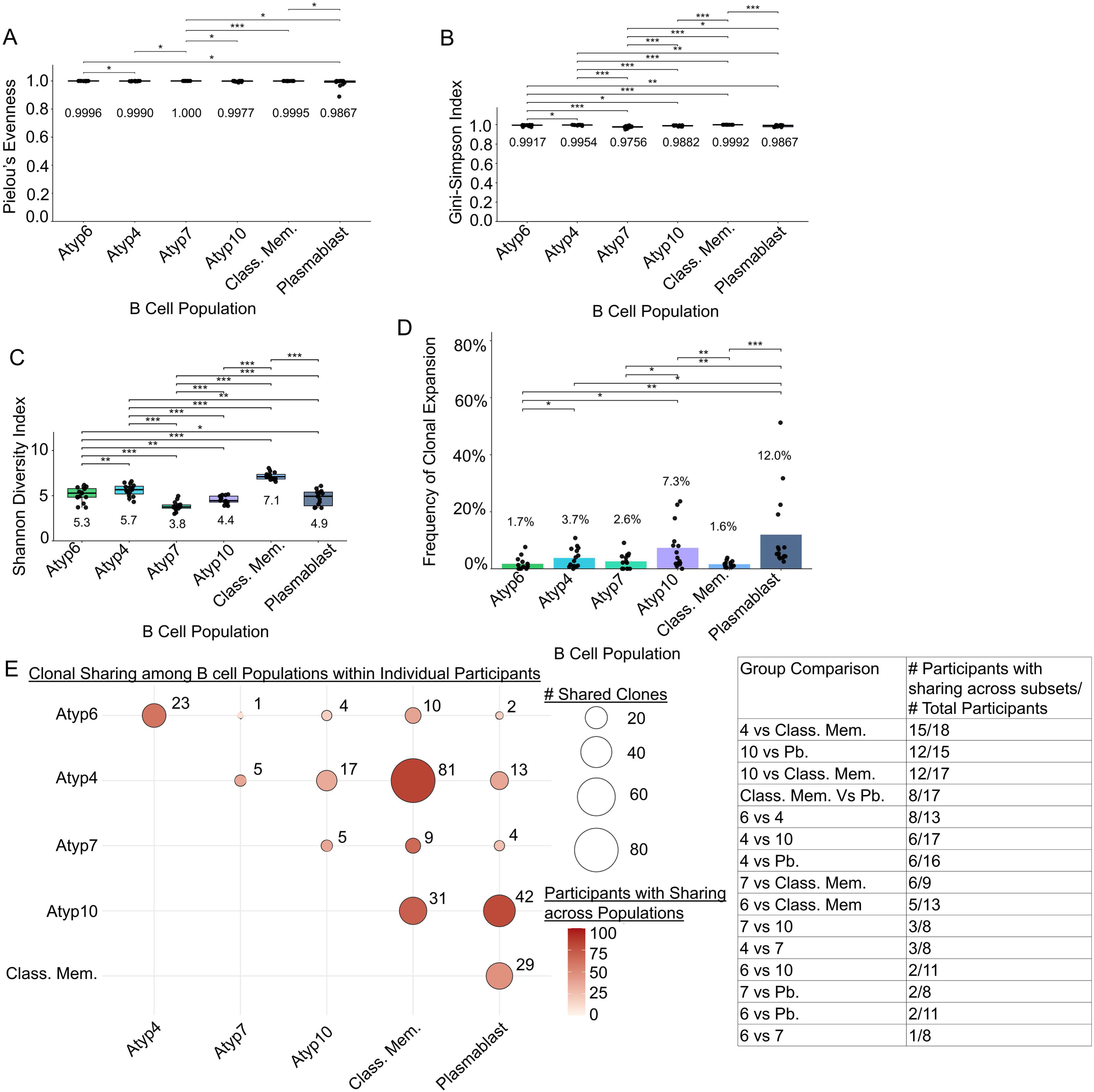
Atypical B cell populations are clonally diverse, with clonal overlap identified across atypical, classical memory, and plasmablast B cell populations within an individual. B cells were isolated and transcriptional and immune repertoire profiling was performed as in Figure 1 and Methods. **A)** Pielou’s evenness, **B)** Gini Simpson index, **C)** Shannon diversity index and D) frequency of clonal expansion were calculated as in Methods across atypical, classical memory, and plasmablast B cell populations based on clonotype IDs, assigned as in Methods. Individuals with n < 20 cells in a given B cell population were excluded from the diversity and clonal expansion analyses. Individual participants (n = 15) are plotted as points and means are listed for each subset. Cells per cluster are listed: Cluster 6 (n = 3,249), Cluster 4 (n = 5,088), Cluster 7 (n = 785), Cluster 10 (n = 1,611), Class. Mem. (n = 21,169), and Plasmablast (n = 2,987). **E)** Circles indicate shared clones identified in the two B cell populations that intersect at the circle, with size indicating the total number of shared clonotypes summed across n = 22 total participants (value listed next to circle). Shading of the circle indicates the frequency of participants that contributed to the total number of shared clones. Cell populations with n < 20 cells, singlets, and any individuals that did not have at least two subsets that met these criteria were excluded from analysis. For panels A-D, Freidman test with paired Wilcoxon signed-rank test (Benjamin Hochberg correction), **** p < 0.0001, *** p < 0.001, ** p < 0.01, * p < 0.05, all other comparisons are not significant.

We next asked whether shared clones exist across atypical, classical memory, and plasmablast B cell populations within individual participants. Atypical Cluster 4 and classical memory B cells exhibited the highest number of shared clones within participants (n = 81 total shared clones in 15/18 participants), while Atypical cluster 7 shared n = 42 clones (12/15 participants) with the plasmablast subset (**Figure 7E**). Clonal sharing was observed across atypical B clusters, with Cluster 4 and Cluster 6 sharing n=23 clones (**Figure 7E**). Individual participant level clonal sharing is shown in **Supplemental Figure S12B.** Shared clones between atypical and plasmablast subsets align with prior reports indicating that atypical B cells can differentiate into antibody secreting cells upon stimulation (18, 26).

### 3.10 Insulin-binding B cells are found at an increased frequency in Atypical Cluster 10 relative to other atypical or classical memory clusters

We previously identified insulin binding B cells in a clonally expanded CD27+ CD21lo atypical B cell subset in Stage 1 T1D individuals (27); therefore, we recombinantly expressed BCRs as monoclonal antibody (mAb) to determine if insulin specificities exist among the four atypical B cell subsets identified here. BCRs were prioritized for recombinant expression if they were transcriptionally defined as an atypical B cell cluster and were clonally expanded. Within a given clonotype, a clone was prioritized for expression if it was mutated and/or class switched. BCRs from classical memory and plasmablast clusters were included for comparison. BCRs from n=143 expanded clones and n=17 non-expanded clones were designed and recombinantly expressed as mAb, as in Methods (**Supplemental Table S10**) and screened for binding to insulin by ELISA (**Figure 8A**). Mouse mAb125, which supports T1D as a BCR in NOD mice (51), was modified to express a human IgG1 Fc for use as an anti-insulin IgG positive control (humAb125) that represents a pathological anti-insulin BCR benchmark, as in (27). The positive cutoff for initial screening was O.D. > 3-fold above the IgG negative control (O.D. = 0.33) (**Figure 8A**). Antibodies that met the initial insulin binding cutoff were rescreened to confirm insulin binding across a range of mAb concentrations (**Figure 8B**); 8.6% of Classical Memory mAbs (5/58), 5.7% of Atypical Cluster 4 mAbs (2/35), 31.3% of Atypical Cluster 10 mAbs (5/16), and 10.5% of Plasmablast mAbs (2/19) bound insulin by ELISA (**Figure 8C**).

**Figure 8.**
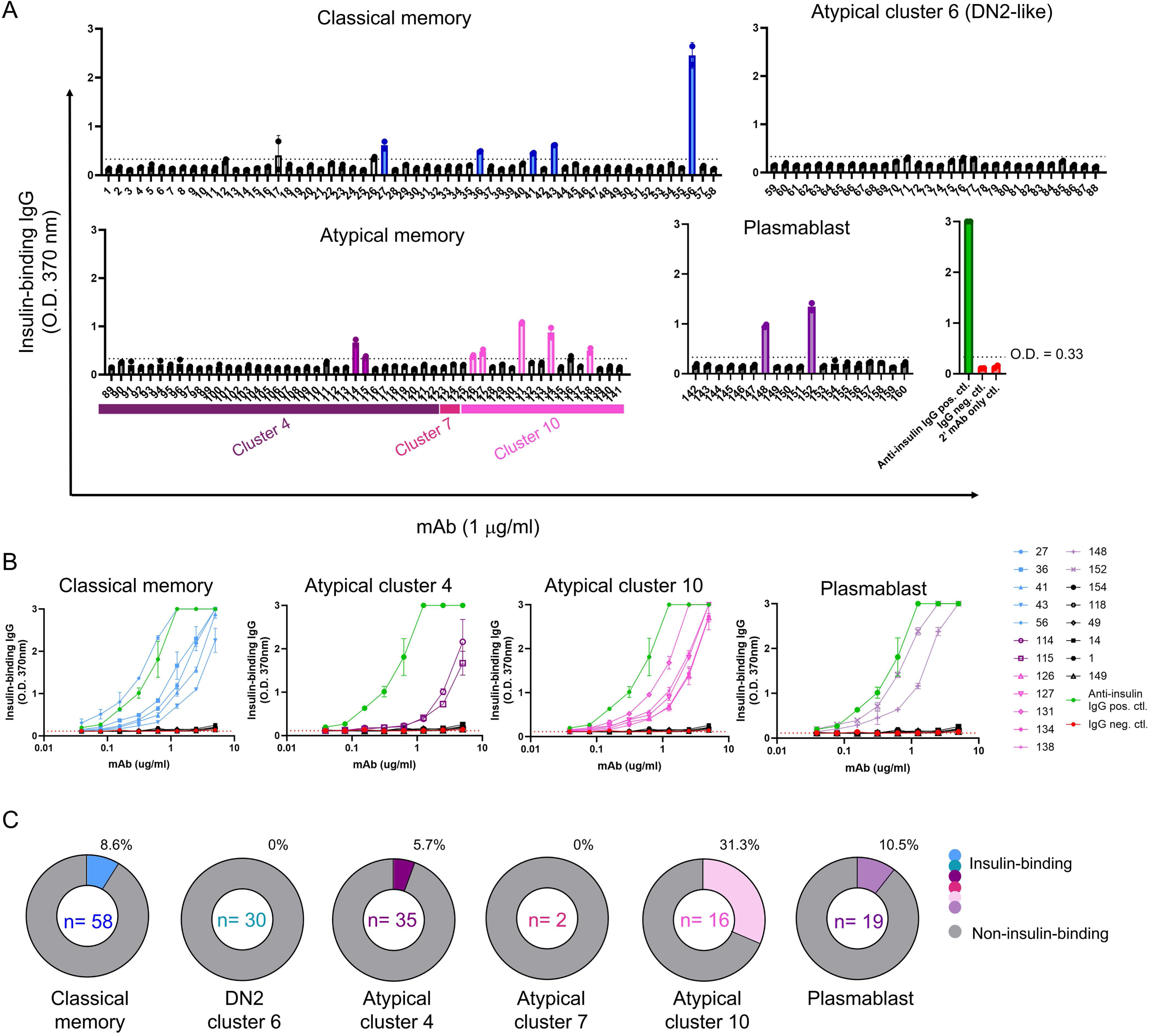
Insulin-binding B cells are identified in Atypical Clusters 4 and 10. BCRs were sequenced by single-cell BCR-seq as in Methods. BCRs from the classical memory, atypical memory clusters 4, 6 (DN2-like), 7, and 10, and plasmablast subsets were prioritized for recombinant expression as IgG1 mAbs based on the following criteria: clonally expanded, mutated, and class-switched. **A)** Recombinant mAbs (n = 160) identified from n = 12 early-stage T1D participants were screened at 1 mg/ml for insulin autoreactivity by insulin-coated ELISA. Anti-insulin mouse BCR (mAb 125) expressed as human IgG1 (humAb125) was designed as a positive control, as the murine version of this mAb supports diabetes development in NOD mice (51). Anti-viral mAb 2526 was used as a negative control (70). mAbs were classified as insulin-binding if both technical replicates exhibited an O.D. > 0.33 (3-fold > IgG neg. ctl.). **B)** mAbs that screened positive in panel A were serially diluted 1:2 from 5 mg/ml and compared to controls as in *(A)*, with insulin ELISA IgG binding curves broken down by B cell subset. **C)** The frequency of anti-insulin mAbs in each B cell population is shown, labeled outside the pie chart, with the total number of mAbs screened in the center.

The anti-insulin mAbs were isolated from n = 6 T1D individuals, and 64% of anti-insulin mAbs (9/14) exhibited the CD27+ CD21lo phenotype (P5), while 36% of anti-insulin mAbs (5/14) exhibited the CD27+ CD21+ phenotype (P6) (**Supplemental Table S11**). Anti-insulin mAb isotype and V gene usage ranged; IGKV3-11 was most abundant and paired with different VH genes (**Supplemental Table S11**). VH mutation ranged from 0.35% - 16.14% and VL mutation ranged from 0% – 8.96% in the anti-insulin mAbs identified here (**Supplemental Table S11**). Overall, insulin-binding B cells were detected among both atypical and classical memory populations, with the highest relative enrichment noted in Atypical Cluster 10, which had the highest proportion of clonal expansion and isotype switching amongst the four atypical B cell populations profiled here.

## 4 Discussion

Atypical B cells are of increasing interest in chronic infection and autoimmunity. Leaders in the field recently highlighted the need to clearly define atypical B cell populations by features including surface markers, transcriptional programming, somatic hypermutation, class switching, location, and the potential to differentiate into antibody secreting cells (15, 16). To address this knowledge gap in human T1D, we designed a strategy to enrich for atypical B cells, in addition to classical memory and plasmablasts, to provide increased depth and power to identify previously unrecognized atypical B cell transcriptional and phenotypic heterogeneity over studies that more broadly focus on total B cells. Examination of clonal expansion, isotype switching and somatic hypermutation further revealed differences in functional consequences across these populations that naturally arose in the physiologic milieu of the individual. Here, we define four transcriptionally defined atypical B cell populations in early-stage T1D individuals further stratified based on phenotypic differences in key B cell markers. The frequency of isotype switching varied widely across these four atypical B cell clusters, with further heterogeneity in switching observed across phenotypic subsets within a given transcriptionally defined atypical B cell cluster. Similarly, we observed differences in the extent of somatic hypermutation across clusters, and even across phenotypes within clusters, indicating biologic differences in B cell activation and selection within these compartments. Finally, certain atypical B cell populations were enriched for insulin-binding B cells, emphasizing compartments that warrant attention in larger studies, ideally with emphasis on insulin and other islet-reactive B cells, and potentially in future immunotherapy trials.

To emphasize shared vs. unique features of atypical B cell populations, we performed comparative analyses anchored against all classical memory B cells grouped together, since this is another antigen-experienced population that undergoes isotype switch and somatic hypermutation. Heterogeneity across atypical B cell populations is demonstrated by identification of only a single shared gene, FCRL5, upregulated among all atypical B cell populations profiled here compared to classical memory B cells. FCRL5 is one of multiple genes used to identify atypical B cells in humans, as reviewed previously (15, 16). Overexpression of FCRL5 in mice overcame B cell anergy and exacerbated SLE (52), highlighting a potential role for FCRL5 in autoimmunity. Although FCRL5 was upregulated in all atypical B cell populations compared to classical memory B cells, the levels of FCRL5 expression differed across atypical B cell populations in T1D individuals. This emphasized a need to examine transcriptional and phenotypic atypical B cell populations more deeply, which revealed gene expression changes, as well as functional changes (isotype switching and somatic hypermutation), which we detail below.

As summarized in **Figure 9**, Atypical Cluster 6 exhibited the highest expression of FCRL5 and other atypical B cell-defining genes ITGAX, FGR, and SOX5, but downregulated classical memory genes CD27, AIM2, and TNFRSF13B compared to classical memory B cells, with lower average expression of these markers relative to the other three atypical B cell clusters. Cluster 6 upregulated transcription factors TBX21, ZEB2, POU2F2, and LITAF, all shown to be upregulated in atypical B cells compared to other B cell subsets in SLE (41) and malaria infection (37). Cluster 6 contained the greatest proportion of surface CD27-CD21lo CD24-CD38-DN2-like cells, but it also exhibited the most phenotypic diversity (in terms of sorted population abundance). Cluster 6 exhibited increased IgG3 class switching, associated with IFN-γ exposure (53)) and Tbet expression (54), with upregulation of the IgG3-associated gene, DAPP1 (55) compared to the other atypical B cell clusters. Cluster 6 exhibited the lowest levels of somatic hypermutation among atypical and classical B cells, and showed decreased clonal diversity compared to classical memory, as was observed in non-class switched atypical B cells in malaria infection (37). This is consistent with a primary/earlier stage of immune response in which clonal divergence, affinity maturation/selection and isotype switching would be more limited. Cluster 6 showed enriched gene usage of autoreactive-prone IGHV4-34 (56–60), yet we did not find anti-insulin specificities among the Cluster 6 mAbs tested. Given the relatively small number of insulin-binding mAbs profiled (and their expected rarity in the repertoire), this may represent sampling depth limitations, rather than true absence, in the insulin-binding B cell repertoire in humans. Gene skewing, including biased IGHV4-34 usage in one atypical B cell population compared to other atypical B cells, was observed in malaria infection (37). VH4-34 has a longer (germline-encoded) CDR2 loops enriched in Gly/Ser residues and a hydrophobic patch in the framework which might impact binding promiscuity to a variety of antigens (61, 62).

**Figure 9.**
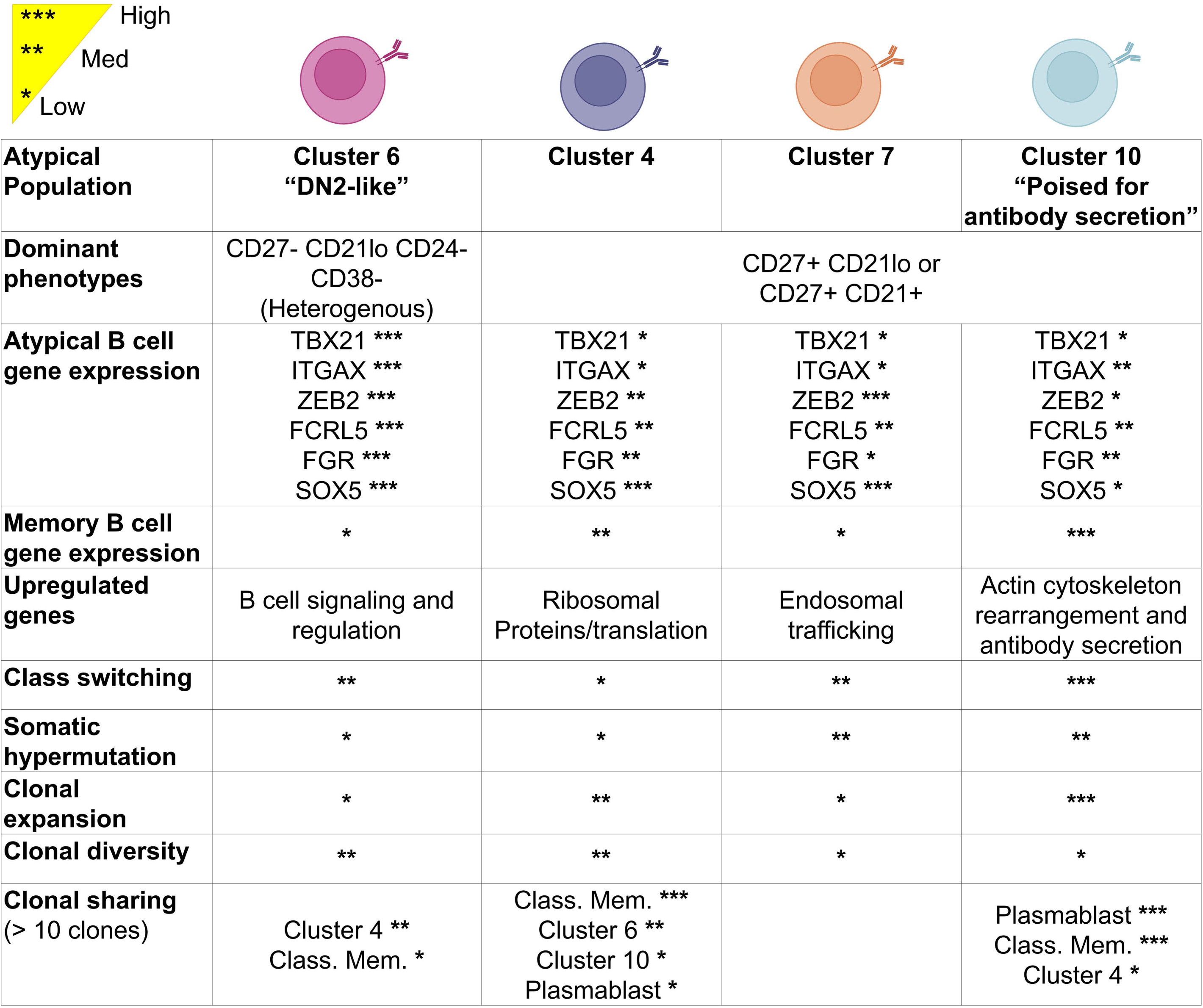
Summary of atypical B cell population heterogeneity in early-stage T1D individuals. Graphical summary outlining differences in phenotype, gene expression, and BCR isotype switching, somatic hypermutation, and clonal expansion among four distinct atypical B cell populations identified in the peripheral blood of early-stage T1D individuals.

Atypical Cluster 4 exhibited increased expression of atypical-defining genes, but maintained CD27 gene expression, as aligned with the most abundant sorted phenotypic populations CD27+CD21lo (P5) and CD27+ CD21+ (P6) in this cluster. Cluster 4 upregulated many ribosomal genes (associated with increased translational output) compared to the other atypical B cell clusters. Ribosomal genes were also upregulated in classical memory B cells compared to Atypical Clusters 6 and 7. Reyes et al. similarly identified atypical B cell populations that upregulated genes encoding ribosomal proteins in malaria infection, HIV infection, and autoimmune SLE (63). Atypical Cluster 4 exhibited the most clonal sharing with classical memory B cells within individuals and exhibited the lowest number of differentially expressed genes compared to classical memory. These findings raise the possibility that Atypical Cluster 4 share similarities in differentiation path with classical memory, as noted in malaria infection studies (64). Anti-insulin specificities were identified in 5.7% of the Atypical Cluster 4 mAbs tested, compared to 8.6% of the classical memory mAbs tested, emphasizing their enrichment in classical and atypical memory compartments. This tracks with our prior high-throughput screening studies, that drove antibody-secretion in vitro to use antibody as a proxy for BCR specificity, which showed increased insulin-binding B cell detection in class-switched, CD27- and CD27+ populations relative to naïve B cells (25).

Atypical Cluster 7 exhibited increased expression of atypical B cell-defining genes, ZEB2 and SOX5, and decreased expression of classical memory genes, CD27 and TNFRSF13B, compared to Atypical Cluster 4 and Cluster 10. Atypical Cluster 7 is the most transcriptionally distinct from classical memory B cells, based on the number of DEGs. Metabolic pathways and antigen processing/presentation pathways were downregulated, while BCR signaling was increased, in Cluster 7 compared to classical memory. Cluster 7 had the lowest Shannon diversity and Gini-Simpson index compared to the other atypical or classical memory B cell populations, yet had a Pielou’s evenness score of 1.0, suggesting this cluster had lower clonotype diversity but a high degree of evenness. Cluster 7 showed minimal clonal sharing with classical memory B cells within individual participants. Thus, in contrast to Atypical Cluster 4, Atypical Cluster 7 may follow a distinct differentiation pathway.

Atypical Cluster 10 was primarily comprised of phenotypic CD27+ CD21+ and CD27+ CD21lo B cells. Cluster 10 expressed increased levels of atypical gene ITGAX and memory genes CD27, AIM2, and TNFRSF13B compared to other atypical B cell clusters. Cluster 10 also expressed increased levels of secreted IgM/IgA pentamer/dimerization gene, JCHAIN, T cell costimulatory gene, CD86, and transcription factor, ARID3A, compared to the other atypical B cell populations. ARID3A (BRIGHT) expression was also increased in Atypical Cluster 4 compared to classical memory. ARID3A is a DNA-binding protein that participates in a protein complex to upregulate immunoglobulin heavy chain transcription, which supports rapid antibody responses (especially in innate-like B cell populations) and class switching, among other functions (65, 66). In line with this, Atypical Cluster 10 exhibited the highest levels of class-switching, somatic hypermutation, and clonal expansion compared to other atypical and classical memory B cell populations. Increased disease activity in SLE patients was associated with increased numbers of ARID3a expressing B cells (67). An increased number of ARID3a+ B cells was also shown in DN B cells in SLE compared to controls, but not in a small cohort or RA patients (67). Cluster 10 exhibited decreased clonal diversity compared to classical memory B cells. Anti-insulin B cells were identified at the highest frequency (31.3%) in Cluster 10, in accordance with increased frequency of anti-insulin B cells in CD27+ memory B cells compared to naïve B cells (25). Cluster 10 appears poised for differentiation into antibody-secreting cells, given its close proximity to plasmablasts on the transcriptionally defined UMAP. Reyes et al. similarly identified an atypical B cell population in malaria infection with increased CD86 expression, increased somatic hypermutation, and the capacity to secrete antibody in response to T cell activation signals (CD40L, IL-2, and IL-21) in vitro (63). CD27+ B cells are a major source of recall responses and exhibit enhanced differentiation into antibody secreting cells (68). Collectively, these data emphasize CD27+ atypical B cells as a population that may prove especially useful in following autoimmune disease progression and/or immunotherapy response, which will require future studies to confirm.

The limitations of our study are as follows. Our results reflect atypical B cell populations in the periphery. Insulin-binding DN2 B cells have been identified in human pancreatic lymph nodes previously (26), but it remains unknown how the findings here will translate to the pancreas or pancreatic lymph nodes. Distinct transcriptional programming across the atypical B cell populations identified here suggest different functional capacities, supported by our examination of functional outcomes such as isotype switch and somatic hypermutation, but future studies are required to directly evaluate how other functions may be perturbed among these atypical B cell populations. Of note, while this is a limitation, it is important to acknowledge that studies using *in vitro* culture, and even organoid systems, may lack important cell types and other attributes only present in 3-D niches *in vivo*). Thus, while more limited, the isotype switching and somatic hypermutation here represent a fully physiologic system that is without these biases.

As a means to circumvent tissue access limitations in humans, our lab previously identified atypical B cell populations in T1D-prone mice which were enriched insulin binding B cells, particularly in the pancreas (14). Insulin-binding B cells were enriched in the B_ND_2 and DN2 B cell subsets in the pancreatic lymph nodes of T1D individuals compared to non-diabetic controls (26). This CD27-IgD+ IgMlo CD21-CXCR5-activated B_ND_2 B cell subset upregulates surface activation markers CD69+ CD86+ CD95+ CD11c+, expresses atypical B cell genes ITGAX, TBX21, FGR, and FCRL5, and is enriched in insulin reactivity in the blood of young-onset T1D participants compared to participants with later disease onset (26). We did not evaluate all of these phenotypic markers, though increased transcription of many of these genes were noted in specific populations captured by our dataset. Human pancreas studies will be necessary to formally evaluate whether atypical B cell populations are also enriched at the site of autoimmune attack or are enriched for islet-reactive B cells in human T1D.

Sera insulin autoantibody levels correlated with frequency of insulin-binding B_ND_2 B cells in T1D individuals (26). In another autoimmune context, FCRL5 expression in CD11c+ atypical B cells differentiated myasthenia gravis subtypes (acetylcholine receptor-positive vs muscle-specific kinase), while CD27 expression on CD11c+ atypical B cells differentiated muscle-specific kinase myasthenia gravis individuals with or without rituximab treatment (69).Therefore, the findings presented here provide a foundation for future studies to evaluate whether differences in T1D age at onset, autoantibody status, clinical features, and/or response to immune-mediating therapies correlate with differences in the atypical B cell populations defined here, which could guide future biomarker strategies and/or participant stratification for appropriate treatment regimens based on anticipated drug mechanism of action.

In summary, the data presented here provide deeper definitions of distinct atypical B cell populations, many of which vary widely in classic metrics of B cell activation, such as isotype switching and increased BCR mutation, but also in transcriptional programming. T1D is a relatively prevalent disease, for which national and international screening programs exist to detect individuals at early, pre-symptomatic disease stages. From a practical standpoint, such an approach is more difficult to apply to rarer autoimmune diseases, with autoimmune disease patients often receiving clinical diagnoses years or even decades after autoimmune responses initiate tissue attack. This study leverages the power afforded by Type 1 Diabetes TrialNet to focus on this difficult to capture, pre-symptomatic phase of autoimmunity, with studies ongoing to differentiate how T1D stage progression may change B cell transcriptional programing. This T1D-centric study provides an outline for similar studies in other autoimmune datasets to provide insight into shared features of autoimmunity early in disease pathogenesis.

## 6 Author Contributions

Conceptualization: L.E.B., R.H.B. Methodology: L.E.B., L.M.C, B.H., R.H.B, Formal analysis: L.E.B., L.M.C., B.H., R.H.M. Resources: D.J.M., R.H.B., Investigation: L.E.B., R.H.B. Writing and editing: L.E.B., L.M.C, D.J.M., R.H.B; Funding acquisition: L.E.B., L.M.C., R.H.B.

## Acknowledgments

The authors thank the Vanderbilt Health Flow Cytometry core for technical assistance with flow cytometry sorting and VANTAGE core staff for technical assistance and expertise regarding library preparation, single-cell sequencing, and data processing. They thank the Type 1 Diabetes TrialNet Study Group’s Pathway to Prevention Study participants and staff. Non-identifying participant metadata were tracked with a REDCap database, a platform initially developed through the Vanderbilt CTSA (UL1 TR000445). We thank Jon Williams for assistance with Participant metadata RedCap design, and Maggie Padgett for peripheral blood processing. We also thank the University of South Florida Office of Research, Innovation, and Scholarly Endeavors (RISE) for their support of B.H. through their Summer Research Elective program. Cartoons in figures were generated using BioRender. The authors used Claude (Anthropic, Sonnet 5) to edit brief sections of the manuscript for brevity and clarity. The authors subsequently reviewed and edited this content for which they take full responsibility.

## Data Availability

Raw sequencing data are available on the Sequence Read Archive under BioProject accession number PRJNA1489232. VDJ and VJ amino acid sequences for all recombinant mAbs are listed in **Supplemental Table S10**. Sequences for n= 14 anti-insulin monoclonal antibodies characterized in this study have been deposited to GenBank under accession numbers listed in **Supplemental Table S11**.

## Abbreviations

BCR: B cell receptor
CLR: centered-log ratio
DEG: differentially expressed gene
DMSO: dimethyl sulfoxide
DN: double negative
FDR: false discovery rate
FBS: fetal bovine serum
GSEA: gene set enrichment analysis
mAb: monoclonal antibody
NES: normalized enrichment score
NOD: non-obese diabetic
PBMCs: peripheral blood mononuclear cells
PBS: phosphate buffered saline
RA: rheumatoid arthritis
SLE: systemic lupus erythematous
T1D: type 1 diabetes

## References

1. Evans-Molina C, Dor Y, Lernmark A, Mathieu C, Millman JR, Mirmira RG, et al. The Heterogeneity of Type 1 Diabetes: Implications for Pathogenesis, Prevention, and Treatment-2024 Diabetes, Diabetes Care, and Diabetologia Expert Forum. Diabetes. 2025;74(10):1730–47.

2. Kramer KJ, Wilfong EM, Voss K, Barone SM, Shiakolas AR, Raju N, et al. Single-cell profiling of the antigen-specific response to BNT162b2 SARS-CoV-2 RNA vaccine. Nat Commun. 2022;13(1):3466.

3. Pescovitz MD, Greenbaum CJ, Bundy B, Becker DJ, Gitelman SE, Goland R, et al. B-lymphocyte depletion with rituximab and beta-cell function: two-year results. Diabetes Care. 2014;37(2):453–9.

4. Pescovitz MD, Greenbaum CJ, Krause-Steinrauf H, Becker DJ, Gitelman SE, Goland R, et al. Rituximab, B-lymphocyte depletion, and preservation of beta-cell function. N Engl J Med. 2009;361(22):2143–52.

5. Insel RA, Dunne JL, Atkinson MA, Chiang JL, Dabelea D, Gottlieb PA, et al. Staging presymptomatic type 1 diabetes: a scientific statement of JDRF, the Endocrine Society, and the American Diabetes Association. Diabetes Care. 2015;38(10):1964–74.

6. Eisenbarth GS, Gianani R, Yu L, Pietropaolo M, Verge CF, Chase HP, et al. Dual-parameter model for prediction of type I diabetes mellitus. Proc Assoc Am Physicians. 1998;110(2):126–35.

7. Hulbert C, Riseili B, Rojas M, and Thomas JW. B cell specificity contributes to the outcome of diabetes in nonobese diabetic mice. J Immunol. 2001;167(10):5535–8.

8. Serreze DV, Fleming SA, Chapman HD, Richard SD, Leiter EH, and Tisch RM. B lymphocytes are critical antigen-presenting cells for the initiation of T cell-mediated autoimmune diabetes in nonobese diabetic mice. J Immunol. 1998;161(8):3912–8.

9. Silveira PA, Johnson E, Chapman HD, Bui T, Tisch RM, and Serreze DV. The preferential ability of B lymphocytes to act as diabetogenic APC in NOD mice depends on expression of self-antigen-specific immunoglobulin receptors. Eur J Immunol. 2002;32(12):3657–66.

10. Noorchashm H, Lieu YK, Noorchashm N, Rostami SY, Greeley SA, Schlachterman A, et al. I-Ag7-mediated antigen presentation by B lymphocytes is critical in overcoming a checkpoint in T cell tolerance to islet beta cells of nonobese diabetic mice. J Immunol. 1999;163(2):743–50.

11. Serreze DV, Chapman HD, Varnum DS, Hanson MS, Reifsnyder PC, Richard SD, et al. B lymphocytes are essential for the initiation of T cell-mediated autoimmune diabetes: analysis of a new “speed congenic” stock of NOD.Ig mu null mice. J Exp Med. 1996;184(5):2049–53.

12. Kendall PL, Case JB, Sullivan AM, Holderness JS, Wells KS, Liu E, et al. Tolerant anti-insulin B cells are effective APCs. J Immunol. 2013;190(6):2519–26.

13. Wan X, Thomas JW, and Unanue ER. Class-switched anti-insulin antibodies originate from unconventional antigen presentation in multiple lymphoid sites. J Exp Med. 2016;213(6):967–78.

14. Clark LM, McAninch JC, McNitt DH, Padgett ML, Jenkins TW, Bass LE, et al. BCL6 in T cells promotes type 1 diabetes by redirecting fates of insulin-autoreactive B lymphocytes. iScience. 2026;29(6):115990.

15. Sanz I. Human Atypical B Cells. An Overview. Immunol Rev. 2025;334(1):e70058.

16. Eisenbarth SC, Batista F, Cyster J, Elsner R, Kelsoe G, Lund FE, et al. A roadmap for defining “extrafollicular” B cell responses. Immunity. 2025;58(11):2627–45.

17. Rincon-Arevalo H, Wiedemann A, Stefanski AL, Lettau M, Szelinski F, Fuchs S, et al. Deep Phenotyping of CD11c(+) B Cells in Systemic Autoimmunity and Controls. Front Immunol. 2021;12:635615.

18. Jenks SA, Cashman KS, Zumaquero E, Marigorta UM, Patel AV, Wang X, et al. Distinct Effector B Cells Induced by Unregulated Toll-like Receptor 7 Contribute to Pathogenic Responses in Systemic Lupus Erythematosus. Immunity. 2018;49(4):725–39 e6.

19. Nishigori R, Hamatani M, Yoshitomi H, Kimura K, Takata M, Ashida S, et al. CD21lo B-cell subsets are recruited to the central nervous system in acute neuromyelitis optica. Brain. 2025;148(8):2995–3010.

20. Wing E, Sutherland C, Miles K, Gray D, Goodyear CS, Otto TD, et al. Double-negative-2 B cells are the major synovial plasma cell precursor in rheumatoid arthritis. Front Immunol. 2023;14:1241474.

21. Wehr C, Eibel H, Masilamani M, Illges H, Schlesier M, Peter HH, et al. A new CD21low B cell population in the peripheral blood of patients with SLE. Clin Immunol. 2004;113(2):161–71.

22. Wilfong EM, Vowell KN, Bunn KE, Rizzi E, Annapureddy N, Dudenhofer RB, et al. CD19 + CD21(lo/neg) cells are increased in systemic sclerosis-associated interstitial lung disease. Clin Exp Med. 2022;22(2):209–20.

23. Nickerson KM, Smita S, Hoehn KB, Marinov AD, Thomas KB, Kos JT, et al. Age-associated B cells are heterogeneous and dynamic drivers of autoimmunity in mice. J Exp Med. 2023;220(5).

24. Smith MJ, Packard TA, O’Neill SK, Henry Dunand CJ, Huang M, Fitzgerald-Miller L, et al. Loss of anergic B cells in prediabetic and new-onset type 1 diabetic patients. Diabetes. 2015;64(5):1703–12.

25. Joosse BA, Jackson JH, Cisneros A, Santhin AB, Smith SA, Moore DJ, et al. High-Throughput Detection of Autoantigen-Specific B Cells Among Distinct Functional Subsets in Autoimmune Donors. Front Immunol. 2021;12:685718.

26. Stensland ZC, Magera CA, Broncucia H, Gomez BD, Rios-Guzman NM, Wells KL, et al. Identification of an anergic BND cell-derived activated B cell population (BND2) in young-onset type 1 diabetes patients. J Exp Med. 2023;220(8).

27. Bass LE, McDonnell WJ, Brannon CT, Kumar NP, Mallal SA, Georgiev IS, et al. Stage 1 type 1 diabetes memory B lymphocytes transcriptionally differ from healthy controls and harbor insulin-binding specificities. Immunohorizons. 2025;9(11).

28. Steck AK, Johnson K, Barriga KJ, Miao D, Yu L, Hutton JC, et al. Age of islet autoantibody appearance and mean levels of insulin, but not GAD or IA-2 autoantibodies, predict age of diagnosis of type 1 diabetes: diabetes autoimmunity study in the young. Diabetes Care. 2011;34(6):1397–9.

29. Nakayama M, Abiru N, Moriyama H, Babaya N, Liu E, Miao D, et al. Prime role for an insulin epitope in the development of type 1 diabetes in NOD mice. Nature. 2005;435(7039):220–3.

30. Yu L, Robles DT, Abiru N, Kaur P, Rewers M, Kelemen K, et al. Early expression of antiinsulin autoantibodies of humans and the NOD mouse: evidence for early determination of subsequent diabetes. Proc Natl Acad Sci U S A. 2000;97(4):1701–6.

31. Felton JL, Maseda D, Bonami RH, Hulbert C, and Thomas JW. Anti-Insulin B Cells Are Poised for Antigen Presentation in Type 1 Diabetes. J Immunol. 2018;201(3):861–73.

32. Harris PA, Taylor R, Thielke R, Payne J, Gonzalez N, and Conde JG. Research electronic data capture (REDCap)--a metadata-driven methodology and workflow process for providing translational research informatics support. J Biomed Inform. 2009;42(2):377–81.

33. Gupta NT, Vander Heiden JA, Uduman M, Gadala-Maria D, Yaari G, and Kleinstein SH. Change-O: a toolkit for analyzing large-scale B cell immunoglobulin repertoire sequencing data. Bioinformatics. 2015;31(20):3356–8.

34. Alamyar E, Duroux P, Lefranc MP, and Giudicelli V. IMGT((R)) tools for the nucleotide analysis of immunoglobulin (IG) and T cell receptor (TR) V-(D)-J repertoires, polymorphisms, and IG mutations: IMGT/V-QUEST and IMGT/HighV-QUEST for NGS. Methods Mol Biol. 2012;882:569–604.

35. Glass DR, Tsai AG, Oliveria JP, Hartmann FJ, Kimmey SC, Calderon AA, et al. An Integrated Multi-omic Single-Cell Atlas of Human B Cell Identity. Immunity. 2020;53(1):217–32 e5.

36. Stewart A, Ng JC, Wallis G, Tsioligka V, Fraternali F, and Dunn-Walters DK. Single-Cell Transcriptomic Analyses Define Distinct Peripheral B Cell Subsets and Discrete Development Pathways. Front Immunol. 2021;12:602539.

37. Holla P, Dizon B, Ambegaonkar AA, Rogel N, Goldschmidt E, Boddapati AK, et al. Shared transcriptional profiles of atypical B cells suggest common drivers of expansion and function in malaria, HIV, and autoimmunity. Sci Adv. 2021;7(22).

38. Espinoza DA, Le Coz C, Cruz Cabrera E, Romberg N, Bar-Or A, and Li R. Distinct stage-specific transcriptional states of B cells derived from human tonsillar tissue. JCI Insight. 2023;8(7).

39. Wei C, Anolik J, Cappione A, Zheng B, Pugh-Bernard A, Brooks J, et al. A new population of cells lacking expression of CD27 represents a notable component of the B cell memory compartment in systemic lupus erythematosus. J Immunol. 2007;178(10):6624–33.

40. Roberts AD, Davenport TM, Dickey AM, Ahn R, Sochacki KA, and Taraska JW. Structurally distinct endocytic pathways for B cell receptors in B lymphocytes. Mol Biol Cell. 2020;31(25):2826–40.

41. Dai D, Gu S, Han X, Ding H, Jiang Y, Zhang X, et al. The transcription factor ZEB2 drives the formation of age-associated B cells. Science. 2024;383(6681):413–21.

42. Shankar M, Nixon JC, Maier S, Workman J, Farris AD, and Webb CF. Anti-nuclear antibody production and autoimmunity in transgenic mice that overexpress the transcription factor Bright. J Immunol. 2007;178(5):2996–3006.

43. Oldham AL, Miner CA, Wang HC, and Webb CF. The transcription factor Bright plays a role in marginal zone B lymphocyte development and autoantibody production. Mol Immunol. 2011;49(1-2):367–79.

44. Cheng D, Deobagkar-Lele M, Zvezdova E, Choi S, Uehara S, Baup D, et al. Themis2 lowers the threshold for B cell activation during positive selection. Nat Immunol. 2017;18(2):205–13.

45. Nair L, Zhang W, Laffleur B, Jha MK, Lim J, Lee H, et al. Mechanism of noncoding RNA-associated N(6)-methyladenosine recognition by an RNA processing complex during IgH DNA recombination. Mol Cell. 2021;81(19):3949–64 e7.

46. Vidarsson G, Dekkers G, and Rispens T. IgG subclasses and allotypes: from structure to effector functions. Front Immunol. 2014;5:520.

47. Rivera CE, Zhou Y, Chupp DP, Yan H, Fisher AD, Simon R, et al. Intrinsic B cell TLR-BCR linked coengagement induces class-switched, hypermutated, neutralizing antibody responses in absence of T cells. Sci Adv. 2023;9(17):eade8928.

48. Elsner RA, and Shlomchik MJ. Germinal Center and Extrafollicular B Cell Responses in Vaccination, Immunity, and Autoimmunity. Immunity. 2020;53(6):1136–50.

49. Good KL, Avery DT, and Tangye SG. Resting human memory B cells are intrinsically programmed for enhanced survival and responsiveness to diverse stimuli compared to naive B cells. J Immunol. 2009;182(2):890–901.

50. Tangye SG, Avery DT, Deenick EK, and Hodgkin PD. Intrinsic differences in the proliferation of naive and memory human B cells as a mechanism for enhanced secondary immune responses. J Immunol. 2003;170(2):686–94.

51. Acevedo-Suarez CA, Hulbert C, Woodward EJ, and Thomas JW. Uncoupling of anergy from developmental arrest in anti-insulin B cells supports the development of autoimmune diabetes. J Immunol. 2005;174(2):827–33.

52. Ono C, Tanaka S, Myouzen K, Iwasaki T, Ueda M, Oda Y, et al. Upregulated Fcrl5 disrupts B cell anergy and causes autoimmune disease. Front Immunol. 2023;14:1276014.

53. Janeway CA, Jr., Travers P, Walport M, and Shlomchik M. Immunobiology. New York: Garland Science; 2001.

54. Peng SL, Szabo SJ, and Glimcher LH. T-bet regulates IgG class switching and pathogenic autoantibody production. Proc Natl Acad Sci U S A. 2002;99(8):5545–50.

55. Fournier E, Isakoff SJ, Ko K, Cardinale CJ, Inghirami GG, Li Z, et al. The B cell SH2/PH domain-containing adaptor Bam32/DAPP1 is required for T cell-independent II antigen responses. Curr Biol. 2003;13(21):1858–66.

56. Wardemann H, Yurasov S, Schaefer A, Young JW, Meffre E, and Nussenzweig MC. Predominant autoantibody production by early human B cell precursors. Science. 2003;301(5638):1374–7.

57. Thompson KM, Sutherland J, Barden G, Melamed MD, Randen I, Natvig JB, et al. Human monoclonal antibodies against blood group antigens preferentially express a VH4-21 variable region gene-associated epitope. Scand J Immunol. 1991;34(4):509–18.

58. Cappione AJ, Pugh-Bernard AE, Anolik JH, and Sanz I. Lupus IgG VH4.34 antibodies bind to a 220-kDa glycoform of CD45/B220 on the surface of human B lymphocytes. J Immunol. 2004;172(7):4298–307.

59. van Vollenhoven RF, Bieber MM, Powell MJ, Gupta PK, Bhat NM, Richards KL, et al. VH4-34 encoded antibodies in systemic lupus erythematosus: a specific diagnostic marker that correlates with clinical disease characteristics. J Rheumatol. 1999;26(8):1727–33.

60. Bashford-Rogers RJM, Bergamaschi L, McKinney EF, Pombal DC, Mescia F, Lee JC, et al. Analysis of the B cell receptor repertoire in six immune-mediated diseases. Nature. 2019;574(7776):122–6.

61. Langley DB, Jara CJ, Henry JY, Reed JH, Goodnow CC, and Christ D. Framework-mediated binding of foreign and self-glycans by IGHV4-34 antibodies. Front Immunol. 2026;17:1767837.

62. Mockridge CI, Chapman CJ, Spellerberg MB, Sheth B, Fleming TP, Isenberg DA, et al. Sequence analysis of V(4-34)-encoded antibodies from single B cells of two patients with systemic lupus erythematosus (SLE). Clin Exp Immunol. 1998;114(1):129–36.

63. Reyes RA, Batugedara G, Dutta P, Reers AB, Garza R, Ssewanyana I, et al. Atypical B cells consist of subsets with distinct functional profiles. iScience. 2023;26(12):108496.

64. Sutton HJ, Aye R, Idris AH, Vistein R, Nduati E, Kai O, et al. Atypical B cells are part of an alternative lineage of B cells that participates in responses to vaccination and infection in humans. Cell Rep. 2021;34(6):108684.

65. Webb CF, Das C, Eaton S, Calame K, and Tucker PW. Novel protein-DNA interactions associated with increased immunoglobulin transcription in response to antigen plus interleukin-5. Mol Cell Biol. 1991;11(10):5197–205.

66. Herrscher RF, Kaplan MH, Lelsz DL, Das C, Scheuermann R, and Tucker PW. The immunoglobulin heavy-chain matrix-associating regions are bound by Bright: a B cell-specific trans-activator that describes a new DNA-binding protein family. Genes Dev. 1995;9(24):3067–82.

67. Ward JM, Rose K, Montgomery C, Adrianto I, James JA, Merrill JT, et al. Disease activity in systemic lupus erythematosus correlates with expression of the transcription factor AT-rich-interactive domain 3A. Arthritis Rheumatol. 2014;66(12):3404–12.

68. Sanz I, Wei C, Jenks SA, Cashman KS, Tipton C, Woodruff MC, et al. Challenges and Opportunities for Consistent Classification of Human B Cell and Plasma Cell Populations. Front Immunol. 2019;10:2458.

69. Sikorski PM, Kaminski HJ, Vincent A, Bauman T, Jacobson L, and Kusner LL. Subtype-specific atypical B cell profiles in myasthenia gravis reveal distinct immunopathological pathways. Front Immunol. 2025;16:1608160.

