## Supplemental Data for "Transcriptionally distinct atypical B cell populations contribute to insulin autoreactivity and show differential activation, isotype switching, and somatic hypermutation in early-stage type 1 diabetes individuals"

### Supplemental Tables

**Supplemental Table S1. Participant Inclusion/Exclusion Criteria [A]**

| Inclusion Criteria | Exclusion Criteria |
|---|---|
| | Diagnosis of diabetes |
| Between the ages of 2 and 45 and have a first degree relative previously diagnosed with T1D | Previous or current use of medications to control hyperglycemia |
| Between the ages of 2 and 20 and have a second-degree relative previously diagnosed with T1D | Current use of immunosuppressive or immunomodulatory therapies |
| Between the ages of 2 and 45 and have already tested positive for at least one autoantibody | Fever 101°F in preceding 48hrs of clinic visit |
| | Receiving antibiotic therapy at clinic visit |
| | Weight less than 8 kg |
| | Known pregnancy |
| | Anemia at time of study initiation (Hgb <12) |
| | Comorbid conditions which would render giving blood samples a risk |
| | Chronic use of immune suppressive or depleting medications |
| | Prior use of immune modulating medications |
| | Inability to meet study visit requirements |
| | Ongoing use of Accutane |

[A] Individuals were enrolled at Vanderbilt through their participation in Type 1 Diabetes TrialNet Pathway to Prevention

| Supplemental Table S2. Individual participant metadata | | | | | |
|---|---|---|---|---|---|
| **Participant** | **Age (Years)** | **Sex** | **Positive islet autoantibodies [A,B]** | **Glucose tolerance test result [B,C,D]** | **T1D Stage** |
| 1 | 21 | F | MIAA, GAD65, ICA, ZNT8 | Impaired | 2 |
| 2 | 18 | F | GAD65, IA2, ICA, ZNT8 | Impaired | 2 |
| 3 | 21 | M | GAD65, IA2, ICA, ZNT8 | Normal | 1 |
| 4 | 22 | F | MIAA [E] | Normal | 1 |
| 5 | 20 | F | GAD65, IA2, ICA, ZNT8 | Impaired | 2 |
| 6 | 13 | M | MIAA, GAD65, IA2, ICA | Normal | 1 |
| 7 | 8 | M | MIAA, GAD65, IA2, ICA | Impaired | 2 |
| 8 | 7 | F | GAD65, IA2, ZNT8 | Impaired | 2 |
| 9 | 17 | M | MIAA, GAD65 | Impaired | 2 |
| 10 | 3 | F | MIAA, GAD65, IA2, ICA, ZNT8 | Impaired | 2 |
| 11 | 8 | F | MIAA, GAD65, IA2, ICA, ZNT8 | Normal | 1 |
| 12 | 20 | M | MIAA, GAD65, ICA, ZNT8 | Impaired | 2 |
| 13 | 14 | M | MIAA, GAD65, IA2, ICA, ZNT8 | Normal | 1 |
| 14 | 14 | F | MIAA, GAD65 | Impaired | 2 |
| 15 | 7 | F | MIAA, GAD65, IA2, ICA, ZNT8 | Impaired | 2 |
| 16 | 9 | F | MIAA, GAD65, ICA, ZNT8 | Normal | 1 |
| 17 | 17 | F | GAD65, IA2, ICA, ZNT8 | Normal | 1 |
| 18 | 10 | F | MIAA, GAD65, ICA | Normal | 1 |
| 19 | 13 | M | MIAA, GAD65, IA2, ICA, ZNT8 | Normal | 1 |
| 20 | 10 | M | MIAA, GAD65, ICA, ZNT8 | Impaired | 2 |
| 21 | 11 | F | MIAA, GAD65 | Normal | 1 |
| 22 | 6 | M | MIAA, GAD65, IA2, ICA, ZNT8 | Normal | 1 |
| 23 | 6 | F | MIAA, GAD65, IA2, ICA, ZNT8 | Impaired | 2 |
| 24 | 2 | F | MIAA, GAD65, IA2, ICA, ZNT8 | Normal | 1 |

[A] *At initial screening visit, TrialNet Pathway to Prevention study participants are screened for MIAA/GAD65/IA2 autoantibodies only, with positivity triggering additional screening for ICA and ZNT8.*
[B] *Islet autoantibody and oral glucose tolerance test (OGTT) results were obtained at the time of blood draw.*
[C] *Normal OGTT results were defined as follows for blood glucose measurements: fasting < 110 mg/dL, 30, 60, or 90min < 200 mg/dL, and 2h < 140 mg/dL.*
[D] *Impaired OGTT results were defined as follows for blood glucose measurements: fasting ≥ 110 and <126 mg/dL, 30, 60, or 90min ≥ 200 mg/dL, and 2h ≥ 140 and < 200 mg/dL.*
[E] *Participant 4 initially screened positive for ≥ 2 islet autoantibodies but was only single positive for MIAA at blood draw.*

| Supplemental Table S3. Sample multiplexing schema for BDRhapsody single-cell sequencing experiments* | | | |
|---|---|---|---|
| **Experiment 1 (ID: 12462)** | | | |
| Lane 1 | Lane 2 | Lane 3 | Lane 4 |
| Participant 11_P5 | Participant 11_P6 | Participant 11_P8 | Participant 11_P9 |
| Participant 3_P6 | Participant 3_P8 | Participant 3_P9 | Participant 3_P5 |
| Participant 4_P8 | Participant 4_P9 | Participant 4_P5 | Participant 4_P6 |
| Participant 6_P9 | Participant 6_P5 | Participant 6_P6 | Participant 6_P8 |
| Lane 5 | Lane 6 | Lane 7 | Lane 8 |
| Participant 7_P5 | Participant 7_P6 | Participant 7_P8 | Participant 7_P9 |
| Participant 5_P6 | Participant 5_P8 | Participant 5_P9 | Participant 5_P5 |
| Participant 9_P8 | Participant 9_P9 | Participant 9_P5 | Participant 9_P6 |
| Participant 2_P9 | Participant 2_P5 | Participant 2_P6 | Participant 2_P8 |
| **Experiment 2 (ID: 13020)** | | | |
| Lane 1 | Lane 2 | Lane 3 | Lane 4 |
| Participant 13_P5 | Participant 13_P6 | Participant 13_P8 | Participant 13_P9 |
| Participant 16_P6 | Participant 16_P8 | Participant 16_P9 | Participant 16_P5 |
| Participant 19_P8 | Participant 19_P9 | Participant 19_P5 | Participant 19_P6 |
| Participant 21_P9 | Participant 21_P5 | Participant 21_P6 | Participant 21_P8 |
| Lane 5 | Lane 6 | Lane 7 | Lane 8 |
| Participant 12_P5 | Participant 12_P6 | Participant 12_P8 | Participant 12_P9 |
| Participant 14_P6 | Participant 14_P8 | Participant 14_P9 | Participant 14_P5 |
| Participant 15_P8 | Participant 15_P9 | Participant 15_P5 | Participant 15_P6 |
| Participant 20_P9 | Participant 20_P5 | Participant 20_P6 | Participant 20_P8 |
| **Experiment 3 (ID: 13282)** | | | |
| Lane 1 | Lane 2 | Lane 3 | Lane 4 |
| Participant 24_P5 | Participant 24_P6 | Participant 24_P8 | Participant 24_P9 |
| Participant 17_P6 | Participant 17_P8 | Participant 17_P9 | Participant 17_P5 |
| Participant 18_P8 | Participant 18_P9 | Participant 18_P5 | Participant 18_P6 |
| Participant 22_P9 | Participant 22_P5 | Participant 22_P6 | Participant 22_P8 |
| Lane 5 | Lane 6 | Lane 7 | Lane 8 |
| Participant 1_P5 | Participant 1_P6 | Participant 1_P8 | Participant 1_P9 |
| Participant 23_P6 | Participant 23_P8 | Participant 23_P9 | Participant 23_P5 |
| Participant 10_P8 | Participant 10_P9 | Participant 10_P5 | Participant 10_P6 |
| Participant 8_P9 | Participant 8_P5 | Participant 8_P6 | Participant 8_P8 |
| *Supplemental Figure S1 shows a representative flow cytometry sort gating scheme for P5 (CD27+ CD21lo), P6 (CD27+ CD21+), P8 (CD27- CD21lo CD24- CD38-), and P9 (CD27- CD21lo CD24+ CD38+/CD24+ CD38-/CD24- CD38+). | | | |

**Supplement Table S4. Atypical B cell population overlap of DEGs identified in atypical vs. classical memory B cells.**

**[Excel file]**

**Supplement Table S5. List of GO BP B cell pathways screened in gene set enrichment analysis (GSEA).**

**[Excel file]**

**Supplemental Table S6. Upregulated and downregulated gene pathways in atypical B cell populations compared to classical memory B cells.**

**[Excel file]**

**Supplemental Table S7. Unfiltered gene list from differential gene expression analysis across atypical B cell clusters**

**[Excel file]**

| **Supplemental Table S8. B cell transcriptionally and phenotypically defined population cell numbers and downstream analysis exclusion rationale** | | | | | |
|---|---|---|---|---|---|
| Transcriptionally and phenotypically defined population (cluster_sortpop) | Cell number (n) | Downstream analysis exclusion rationale | Transcriptionally and phenotypically defined population (cluster_sortpop) | Cell number (n) | Downstream analysis exclusion rationale |
| 0_P5 | 1222 | | 9_P5 | 75 | Non-B cell |
| 0_P6 | 9538 | | 9_P6 | 207 | Non-B cell |
| 0_P8 | 84 | | 9_P8 | 30 | Non-B cell |
| 0_P9 | 991 | | 9_P9 | 2024 | Non-B cell |
| 1_P5 | 1011 | | 10_P5 | 763 | |
| 1_P6 | 8117 | | 10_P6 | 1257 | |
| 1_P8 | 119 | | 10_P8 | 27 | < 50 cells |
| 1_P9 | 355 | | 10_P9 | 152 | |
| 2_P5 | 272 | | 11_P5 | 1663 | |
| 2_P6 | 688 | | 11_P6 | 230 | |
| 2_P8 | 29 | < 50 cells | 11_P8 | 4 | < 50 cells |
| 2_P9 | 8157 | | 11_P9 | 161 | |
| 3_P5 | 555 | | 12_P5 | 158 | Non-B cell |
| 3_P6 | 6486 | | 12_P6 | 89 | Non-B cell |
| 3_P8 | 25 | < 50 cells | 12_P8 | 38 | Non-B cell |
| 3_P9 | 447 | | 12_P9 | 1365 | Non-B cell |
| 4_P5 | 2628 | | 13_P5 | 30 | < 50 cells |
| 4_P6 | 3526 | | 13_P6 | 338 | |
| 4_P8 | 80 | | 13_P8 | 153 | |
| 4_P9 | 734 | | 13_P9 | 1088 | |
| 5_P5 | 115 | | 14_P5 | 1203 | |
| 5_P6 | 3807 | | 14_P6 | 146 | |
| 5_P8 | 479 | | 14_P8 | 2 | < 50 cells |
| 5_P9 | 2379 | | 14_P9 | 147 | |
| 6_P5 | 1108 | | 15_P5 | 299 | Non-B cell |
| 6_P6 | 193 | | 15_P6 | 1121 | Non-B cell |
| 6_P8 | 1887 | | 15_P8 | 15 | Non-B cell |
| 6_P9 | 1293 | | 15_P9 | 44 | Non-B cell |
| 7_P5 | 940 | | 16_P5 | 50 | |
| 7_P6 | 2661 | | 16_P6 | 244 | |
| 7_P8 | 99 | | 16_P8 | 21 | < 50 cells |
| 7_P9 | 552 | | 16_P9 | 19 | < 50 cells |
| 8_P5 | 489 | | 17_P5 | 40 | < 50 cells |
| 8_P6 | 2297 | | 17_P6 | 194 | |
| 8_P8 | 31 | < 50 cells | 17_P8 | 0 | < 50 cells |
| 8_P9 | 210 | | 17_P9 | 30 | < 50 cells |

**Supplement Table S9. Differentially expressed genes across the four phenotypic sort populations within each transcriptionally defined B cell cluster.**

**[Excel]**

**Supplement Table S10. VH/VL usage and IgH and IgL amino acid sequences of recombinant mAbs tested for insulin reactivity.**

**[Excel]**

| Supplemental Table S11. BCR gene usage and somatic hypermutation of anti-insulin mAbs isolated from Stage 1 T1D individuals. | | | | | | | | | | | | |
|---|---|---|---|---|---|---|---|---|---|---|---|---|
| **mAb ID** | **Seurat Barcode** | **Participant ID** | **Subset** | **Seurat Cluster** | **Phenotypic Population** | **Isotype** | **VH Gene** | **%VH SHM** | **VL Gene** | **%VL SHM** | **GenBank # VH** | **GenBank # VL** |
| 27 | 40913767 | 17 | Class. Mem. | 1 | P6 | IGHD | IGHV3-21 | 16.14 | IGLV3-21 | 8.96 | PZ859007 | PZ859008 |
| 36 | 11128941 | 2 | Class. Mem. | 3 | P6 | IGHM | IGHV3-23 | 6.25 | IGKV3-20 | 4.61 | PZ859025 | PZ859026 |
| 41 | 6968925 | 13 | Class. Mem. | 3 | P6 | IGHA1 | IGHV3-30-3 | 0.35 | IGKV3-11 | 0.72 | PZ859021 | PZ859022 |
| 43 | 44693797 | 2 | Class. Mem. | 3 | P6 | IGHM | IGHV3-74 | 4.86 | IGKV3-11 | 1.08 | PZ859033 | PZ859034 |
| 56 | 16144263 | 11 | Class. Mem. | 8 | P5 | IGHG3 | IGHV3-64D | 4.51 | IGLV2-14 | 1.74 | PZ858989 | PZ858990 |
| 114 | 55304750 | 18 | Atyp4 | 4 | P5 | IGHM | IGHV3-30 | 7.64 | IGLV2-8 | 3.47 | PZ859029 | PZ859030 |
| 115 | 4683919 | 23 | Atyp4 | 4 | P5 | IGHM | IGHV3-74 | 2.78 | IGLV2-11 | 1.39 | PZ859047 | PZ859048 |
| 126 | 53372560 | 11 | Atyp10 | 10 | P5 | IGHA1 | IGHV3-64 | 6.6 | IGKV3-11 | 3.58 | PZ859039 | PZ859040 |
| 127 | 24589754 | 11 | Atyp10 | 10 | P5 | IGHG1 | IGHV1-69 | 6.6 | IGKV2-28 | 0 | PZ859019 | PZ859020 |
| 131 | 35326627 | 11 | Atyp10 | 10 | P5 | IGHD | IGHV4-38-2 | 11.11 | IGKV3-11 | 7.53 | PZ859005 | PZ859006 |
| 134 | 12479747 | 18 | Atyp10 | 10 | P5 | IGHG3 | IGHV3-30 | 5.9 | IGKV3-15 | 2.15 | PZ859009 | PZ859010 |
| 138 | 32216756 | 11 | Atyp10 | 10 | P6 | IGHA1 | IGHV4-59 | 9.12 | IGKV3-11 | 2.51 | PZ859015 | PZ859016 |
| 148 | 7081084 | 11 | Plasmablast | 11 | P5 | IGHA1 | IGHV4-59 | 5.96 | IGKV3-11 | 1.43 | PZ859001 | PZ859002 |
| 152 | 12371031 | 17 | Plasmablast | 11 | P5 | IGHG1 | IGHV1-46 | 10.42 | IGKV1-39 | 7.17 | PZ858997 | PZ858998 |

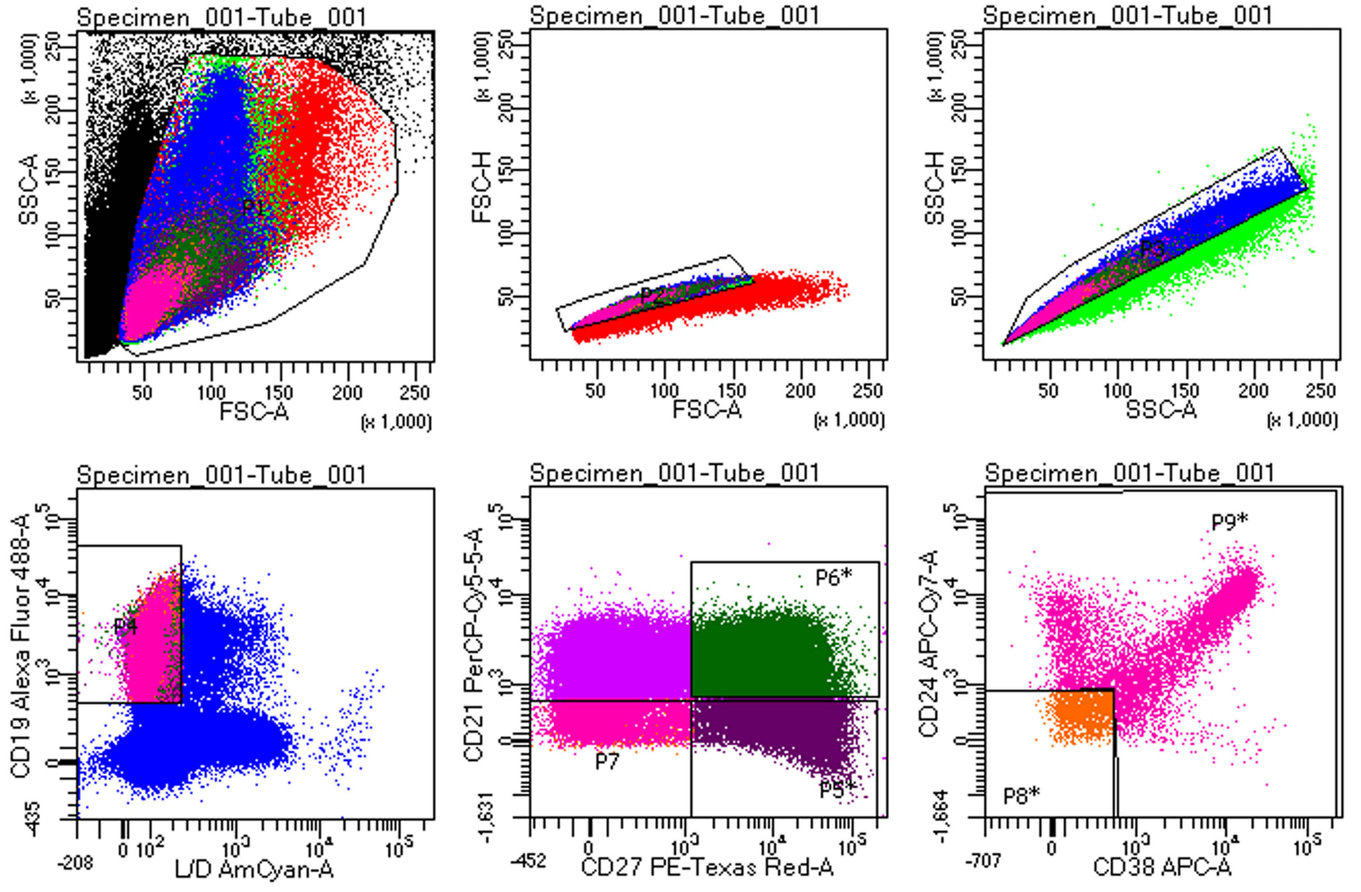

<u>CD19+ live singlet lymphocytes gate:</u>

P5: CD27+ CD21lo

P6: CD27+ CD21+

P8: CD27- CD21lo CD24- CD38-

P9: CD27- CD21lo CD24+ CD38+/CD24- CD38+/ CD24- CD38+

### Supplemental Figures

**Supplemental Figure S1. Representative flow cytometry sorting scheme.** Phenotypic B cell subsets were flow cytometry sorted from n = 24 early-stage T1D individuals' PBMCs, amongst CD19+ live lymphocytes, as follows: (P5) CD27+ CD21lo, (P6) CD27+ CD21+, (P8) CD27- CD21lo CD24- CD38-, and (P9) CD27- CD21lo CD24+ CD38+/CD24+ CD38-/CD24- CD38+, with populations gated as shown in representative plots from one of the participants.

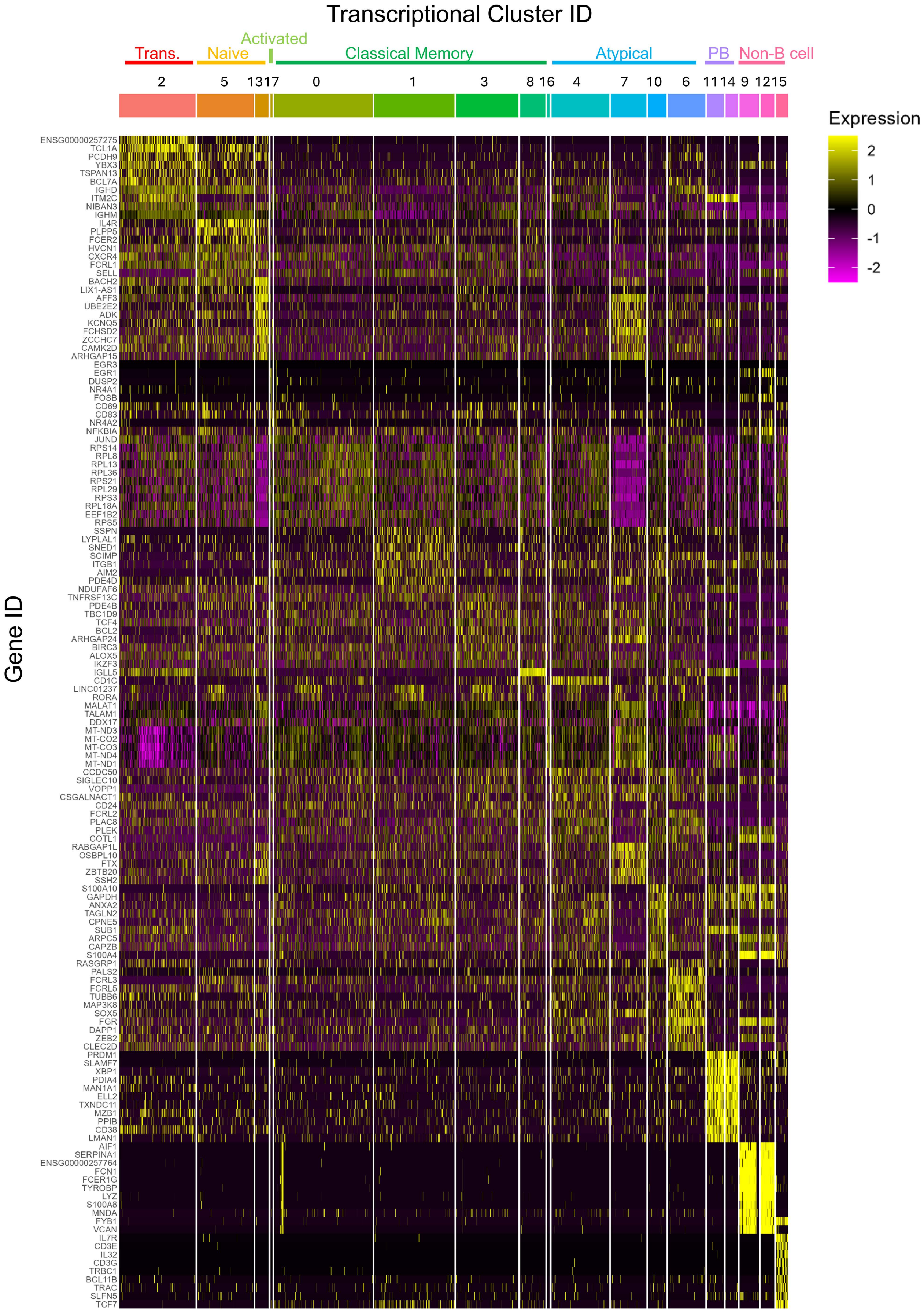

Transcriptional Cluster ID
Trans.
Naive
Activated
Classical Memory
Atypical
PB
Non-B cell
2
5
13
17
0
1
3
8
16
4
7
10
6
11
14
9
12
15
Expression
2
1
0
-1
-2
Gene ID
ENSG00000257275
TCL1A
PCDH9
YBX3
TSPAN13
BCL7A
IGHD
ITM2C
NIBAN3
IGHM
IL4R
PLPP5
FCER2
HVCN1
CXCR4
FCRL1
SELL
BACH2
LIX1-AS1
AFF3
UBE2E2
ADK
KCNQ5
FCHSD2
ZCCHC7
CAMK2D
ARHGAP15
EGR3
EGR1
DUSP2
NR4A1
FOSB
CD69
CD83
NR4A2
NFKBIA
JUND
RPS14
RPL8
RPL13
RPL36
RPS21
RPL29
RPS3
RPL18A
EEF1B2
RPS5
SSPN
LYPLAL1
SNED1
SCIMP
ITGB1
AIM2
PDE4D
NDUFAF6
TNFRSF13C
PDE4B
TBC1D9
TCF4
BCL2
ARHGAP24
BIRC3
ALOX5
IKZF3
IGLL5
CD1C
LINC01237
RORA
MALAT1
TALAM1
DDX17
MT-ND3
MT-CO2
MT-CO3
MT-ND4
MT-ND1
CCDC50
SIGLEC10
VOPP1
CSGALNACT1
CD24
FCRL2
PLAC8
PLEK
COTL1
RABGAP1L
OSBPL10
FTX
ZBTB20
SSH2
S100A10
GAPDH
ANXA2
TAGLN2
CPNE5
SUB1
ARPC5
CAPZB
S100A4
RASGRP1
PALS2
FCRL3
FCRL5
TUBB6
MAP3K8
SOX5
FGR
DAPP1
ZEB2
CLEC2D
PRDM1
SLAMF7
XBP1
PDIA4
MAN1A1
ELL2
TXNDC11
MZB1
PPIB
CD38
LMAN1
AIF1
SERPINA1
ENSG00000257764
FCN1
FCER1G
TYROBP
LYZ
S100A8
MNDA
FYB1
VCAN
IL7R
CD3E
IL32
CD3G
TRBC1
BCL11B
TRAC
SLFN5
TCF7

**Supplemental Figure S2. Transcriptionally defined clusters show differences in gene expression.** B cell multi-omics profiling was performed as in **Figure 1** and Methods. The Seurat FindAllMarkers function was used to identify the top 10 differentially expressed genes in each cluster relative to all other cells in the dataset, using the non-parametric Wilcoxon rank-sum test, adjusted p value $< 0.05$, fold change $>1.5$, expressed by $>30\%$ of the cells in at least one of the comparator groups.

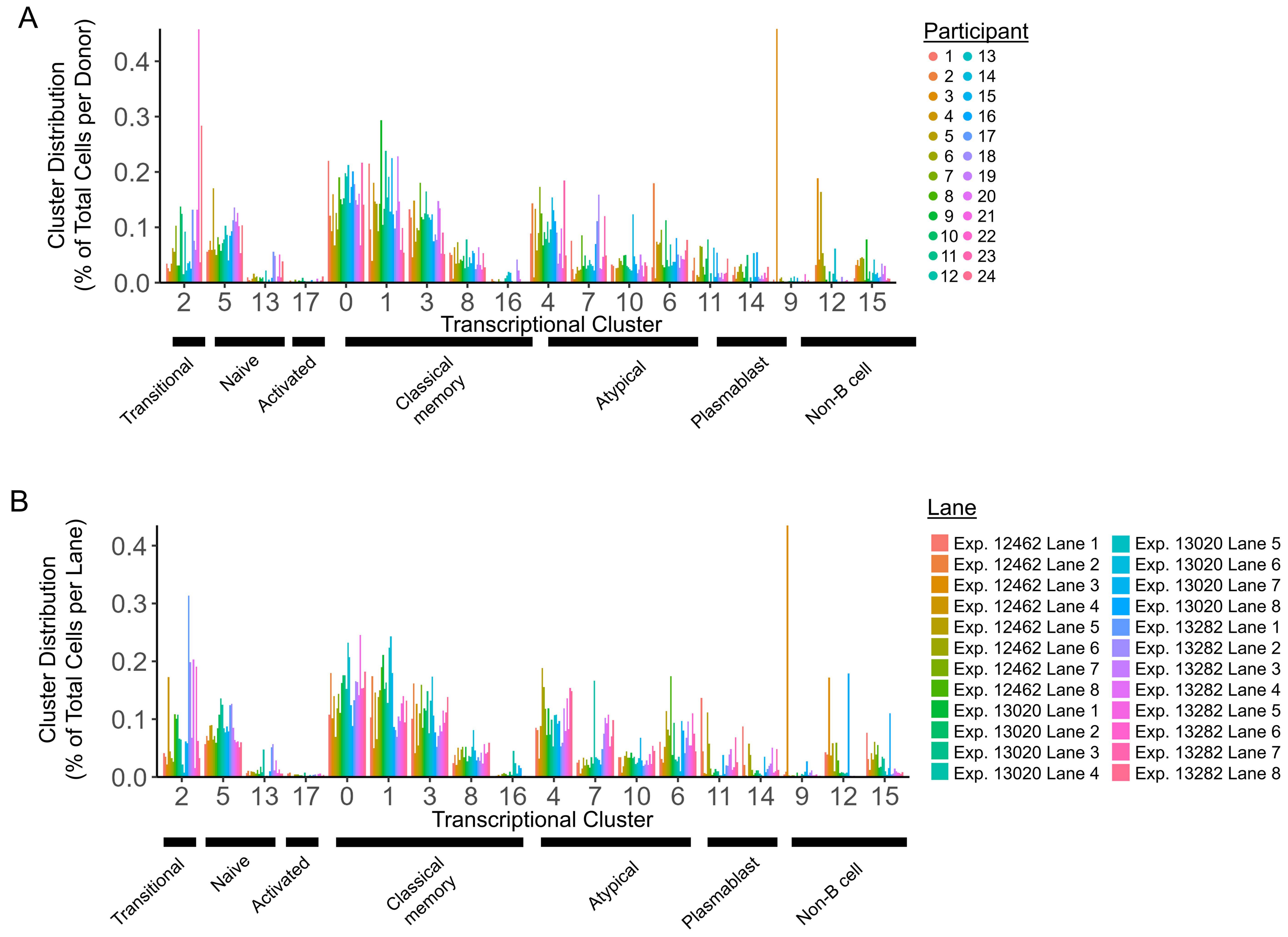
A
Cluster Distribution (% of Total Cells per Donor)
0.0
0.1
0.2
0.3
0.4
2 5 13 17 0 1 3 8 16 4 7 10 6 11 14 9 12 15
Transcriptional Cluster
Transitional
Naive
Activated
Classical memory
Atypical
Plasmablast
Non-B cell
Participant
1 2 3 4 5 6 7 8 9 10 11 12 13 14 15 16 17 18 19 20 21 22 23 24
B
Cluster Distribution (% of Total Cells per Lane)
0.0
0.1
0.2
0.3
0.4
2 5 13 17 0 1 3 8 16 4 7 10 6 11 14 9 12 15
Transcriptional Cluster
Transitional
Naive
Activated
Classical memory
Atypical
Plasmablast
Non-B cell
Lane
Exp. 12462 Lane 1
Exp. 12462 Lane 2
Exp. 12462 Lane 3
Exp. 12462 Lane 4
Exp. 12462 Lane 5
Exp. 12462 Lane 6
Exp. 12462 Lane 7
Exp. 12462 Lane 8
Exp. 13020 Lane 1
Exp. 13020 Lane 2
Exp. 13020 Lane 3
Exp. 13020 Lane 4
Exp. 13020 Lane 5
Exp. 13020 Lane 6
Exp. 13020 Lane 7
Exp. 13020 Lane 8
Exp. 13282 Lane 1
Exp. 13282 Lane 2
Exp. 13282 Lane 3
Exp. 13282 Lane 4
Exp. 13282 Lane 5
Exp. 13282 Lane 6
Exp. 13282 Lane 7
Exp. 13282 Lane 8

**Supplemental Figure S3. Individual participants and BD Rhapsody sequencing lanes are represented across transcriptionally defined B cell clusters.** B cell multi-omics profiling was performed as in **Figure 1** and Methods. Transcriptionally defined clusters were identified as in **Figure 1** and the cluster distribution is shown within n = 24 **A)** individual participants or **B)** BD Rhapsody sequencing lanes (in which four phenotypic populations from different individuals were multiplexed).

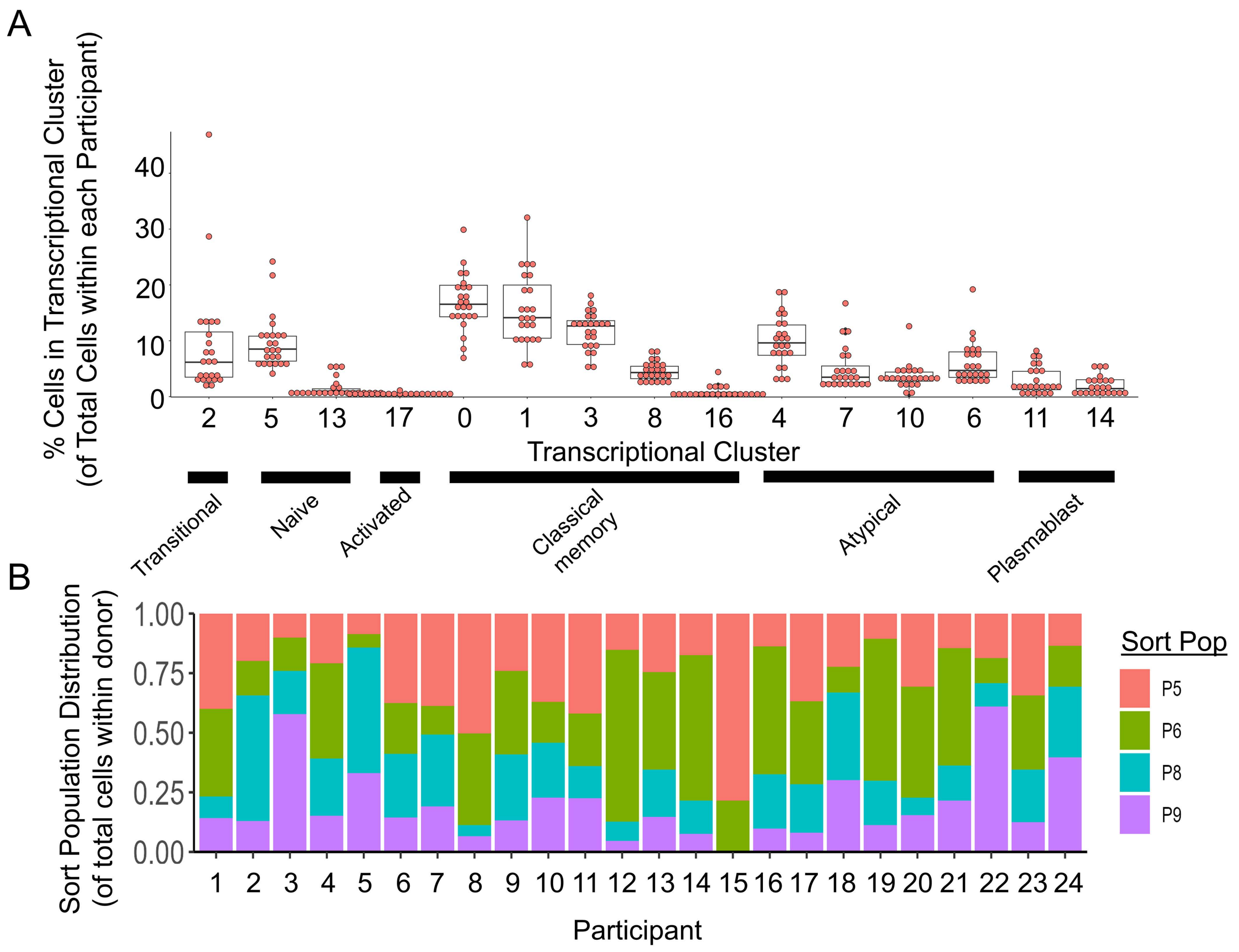
A
% Cells in Transcriptional Cluster
(of Total Cells within each Participant)
0
10
20
30
40
2
5
13
17
0
1
3
8
16
4
7
10
6
11
14
Transcriptional Cluster
Transitional
Naive
Activated
Classical memory
Atypical
Plasmablast
B
Sort Population Distribution
(of total cells within donor)
0.00
0.25
0.50
0.75
1.00
1 2 3 4 5 6 7 8 9 10 11 12 13 14 15 16 17 18 19 20 21 22 23 24
Participant
Sort Pop
P5
P6
P8
P9

**Supplemental Figure S4. Individual participant-level transcriptionally defined cluster frequencies and phenotypic sort population distribution. A)** Transcriptionally defined clusters were identified as in **Figure 1** and the cluster distribution of total cells within each participant is shown. **B)** Phenotypic populations were sorted as in **Figure 1A** and **Supplemental Figure S1** and the distribution across these populations within each of n = 24 individual participants is shown.

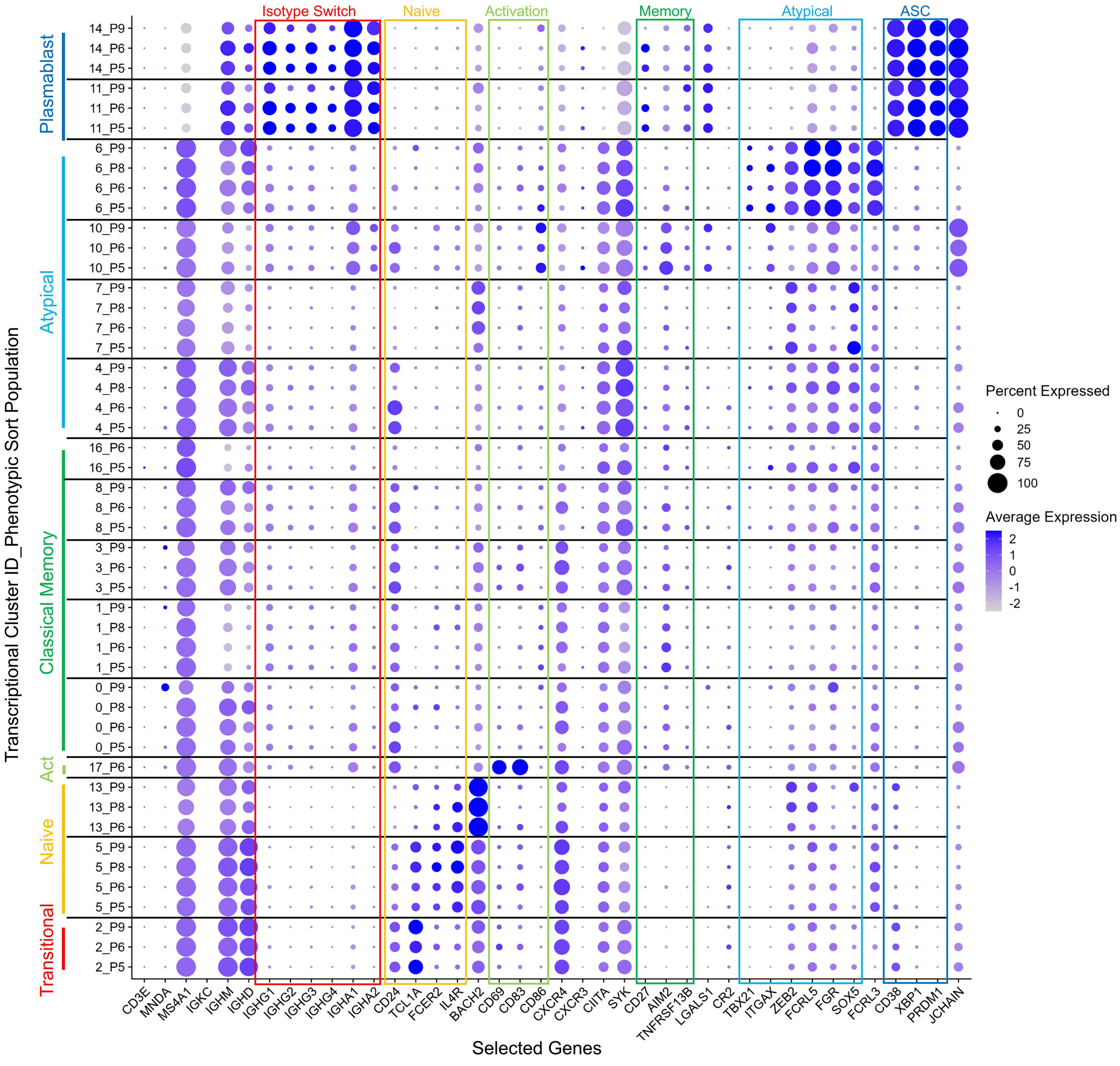

Isotype Switch
Naive
Activation
Memory
Atypical
ASC
Plasmablast
Atypical
Classical Memory
Act
Naive
Transitional
14_P9
14_P6
14_P5
11_P9
11_P6
11_P5
6_P9
6_P8
6_P6
6_P5
10_P9
10_P6
10_P5
7_P9
7_P8
7_P6
7_P5
4_P9
4_P8
4_P6
4_P5
16_P6
16_P5
8_P9
8_P6
8_P5
3_P9
3_P6
3_P5
1_P9
1_P8
1_P6
1_P5
0_P9
0_P8
0_P6
0_P5
17_P6
13_P9
13_P8
13_P6
5_P9
5_P8
5_P6
5_P5
2_P9
2_P6
2_P5
Transcriptional Cluster ID_Phenotypic Sort Population
CD3E
MNDA
MS4A1
IGKC
IGHM
IGHD
IGHG1
IGHG2
IGHG3
IGHG4
IGHA1
IGHA2
CD24
TCL1A
FCER2
IL4R
BACH2
CD69
CD83
CD86
CXCR4
CXCR3
CIITA
SYK
CD27
AIM2
TNFRSF13B
LGALS1
CR2
TBX21
ITGAX
ZEB2
FCRL5
FGR
SOX5
FCRL3
CD38
XBP1
PRDM1
JCHAIN
Selected Genes
Percent Expressed
0
25
50
75
100
Average Expression
2
1
0
-1
-2

**Supplemental Figure S5. Some phenotypic populations within transcriptionally defined atypical B cell clusters show differences in gene expression**. Four distinct phenotypic populations were flow cytometry sorted as in **Figure 1A** and **Supplemental Figure S1** and identified within transcriptionally defined B cell clusters (as depicted in **Figure 1B, D**). Percent and average expression of selected genes (grouped as indicated by colored boxes) are shown for each phenotypically and transcriptionally defined B cell population; populations with < 50 cells were excluded from analysis.

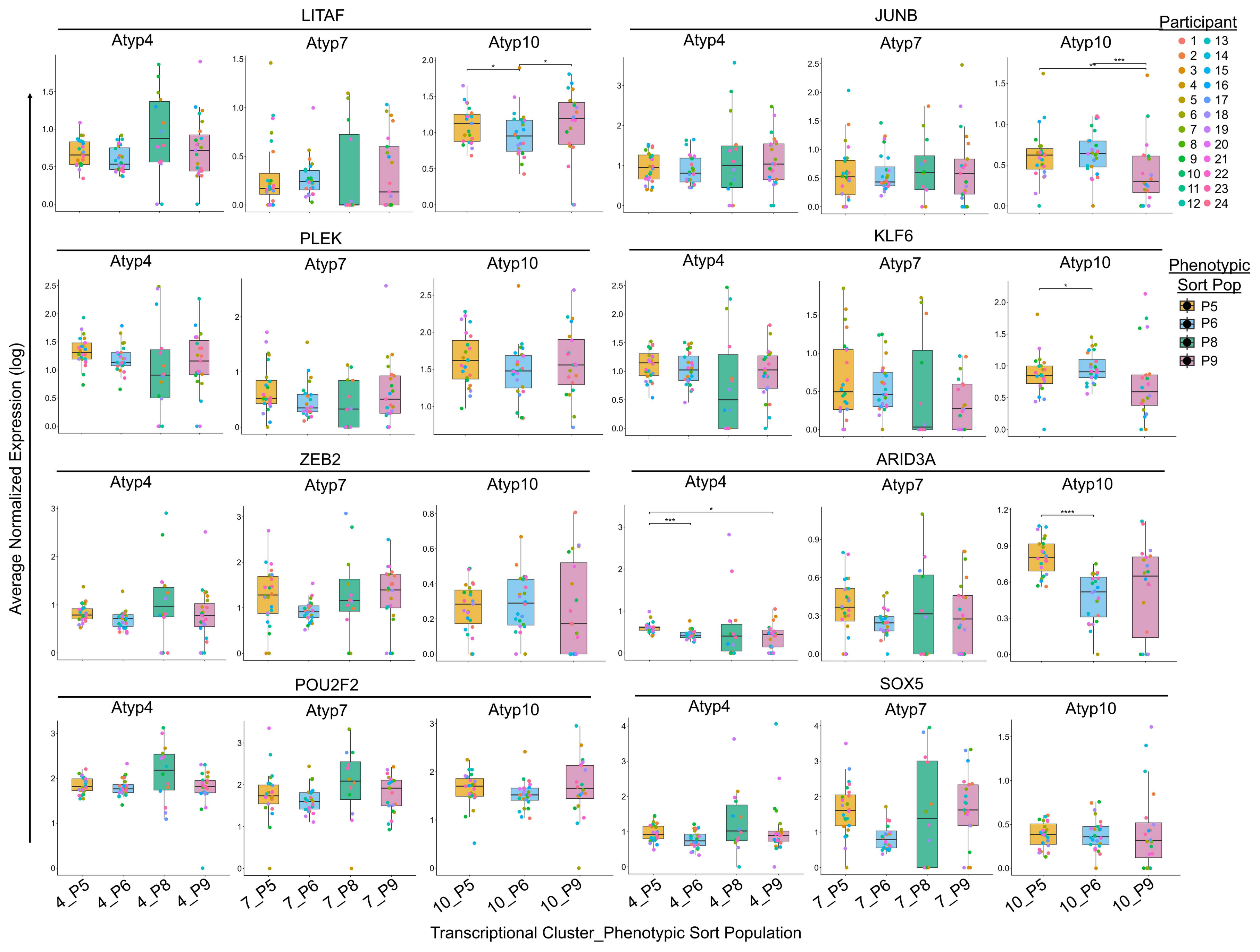

LITAF
JUNB
PLEK
KLF6
ZEB2
ARID3A
POU2F2
SOX5
Atyp4
Atyp7
Atyp10
Participant
Phenotypic Sort Pop
P5
P6
P8
P9
Average Normalized Expression (log)
Transcriptional Cluster_Phenotypic Sort Population
4_P5
4_P6
4_P8
4_P9
7_P5
7_P6
7_P8
7_P9
10_P5
10_P6
10_P9

**Supplemental Figures S6. Phenotypic populations within Atypical Cluster 10 show differential expression of LITAF, JUNB, KLF6, and ARID3A.** B cell multi-omics profiling was performed as in **Figure 1** and Methods. Atypical B cell clusters were subclassified into one of four phenotypically defined sort populations and evaluated for expression of the n = 27 transcription factor genes assayed in **Figure 4B**. Individual participants (n = 24) are plotted as points. Paired statistical analysis was restricted to the number of participants indicated who had cells in all four sort populations: cluster 4 (n = 14), cluster 7 (n = 12), cluster 10 (n = 19). Freidman test with paired Wilcoxon signed-rank test (Benjamin Hochberg correction), **** $p < 0.0001$, *** $p < 0.001$, ** $p < 0.01$, * $p < 0.05$, all other comparisons are not significant.

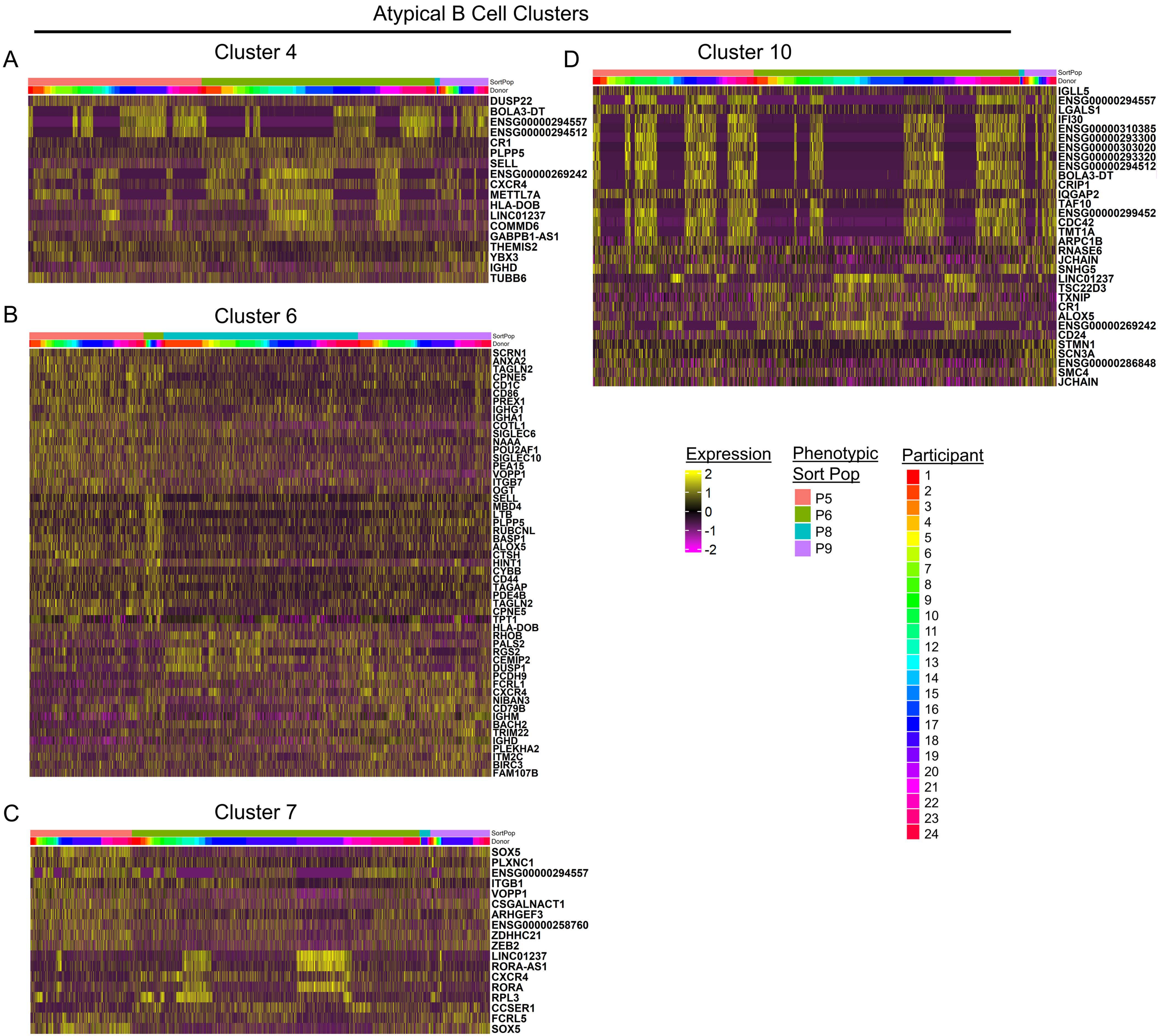

Atypical B Cell Clusters
A
Cluster 4
SortPop
Donor
DUSP22
BOLA3-DT
ENSG00000294557
ENSG00000294512
CR1
PLPP5
SELL
ENSG00000269242
CXCR4
METTL7A
HLA-DOB
LINC01237
COMMD6
GABPB1-AS1
THEMIS2
YBX3
IGHD
TUBB6
B
Cluster 6
SortPop
Donor
SCRN1
ANXA2
TAGLN2
CPNE5
CD1C
CD86
PREX1
IGHG1
IGHA1
COTL1
SIGLEC6
NAAA
POU2AF1
SIGLEC10
PEA15
VOPP1
ITGB7
OGT
SELL
MBD4
LTB
PLPP5
RUBCNL
BASP1
ALOX5
CTSH
HINT1
CYBB
CD44
TAGAP
PDE4B
TAGLN2
CPNE5
TPT1
HLA-DOB
RHOB
PALS2
RGS2
CEMIP2
DUSP1
PCDH9
FCRL1
CXCR4
NIBAN3
CD79B
IGHM
BACH2
TRIM22
IGHD
PLEKHA2
ITM2C
BIRC3
FAM107B
C
Cluster 7
SortPop
Donor
SOX5
PLXNC1
ENSG00000294557
ITGB1
VOPP1
CSGALNACT1
ARHGEF3
ENSG00000258760
ZDHHC21
ZEB2
LINC01237
RORA-AS1
CXCR4
RORA
RPL3
CCSER1
FCRL5
SOX5
D
Cluster 10
SortPop
Donor
IGLL5
ENSG00000294557
LGALS1
IFI30
ENSG00000310385
ENSG00000293300
ENSG00000303020
ENSG00000293320
ENSG00000294512
BOLA3-DT
CRIP1
IQGAP2
TAF10
ENSG00000299452
CDC42
TMT1A
ARPC1B
RNASE6
JCHAIN
SNHG5
LINC01237
TSC22D3
TXNIP
CR1
ALOX5
ENSG00000269242
CD24
STMN1
SCN3A
ENSG00000286848
SMC4
JCHAIN
Expression
2
1
0
-1
-2
Phenotypic Sort Pop
P5
P6
P8
P9
Participant

**Supplemental Figure S7. Differentially expressed genes were identified across phenotypically defined B cell populations within each atypical B cell cluster.** B cell multi-omics profiling was performed as in **Figure 1** and Methods. The top 20 upregulated genes (padj $< 0.05$, FC $> 1.5$, expressed by >30% of the cells in at least one of the comparator groups, where comparator groups required $n \geq 10$ cells) were identified using the Seurat FindAllMarkers function, with DEGs determined amongst phenotypic populations within each of four transcriptionally defined atypical B cell clusters. Individual participant ID is also indicated on heatmaps for **A)** Cluster 4, **B)** Cluster 6, **C)** Cluster 7, and **D)** Cluster 10.

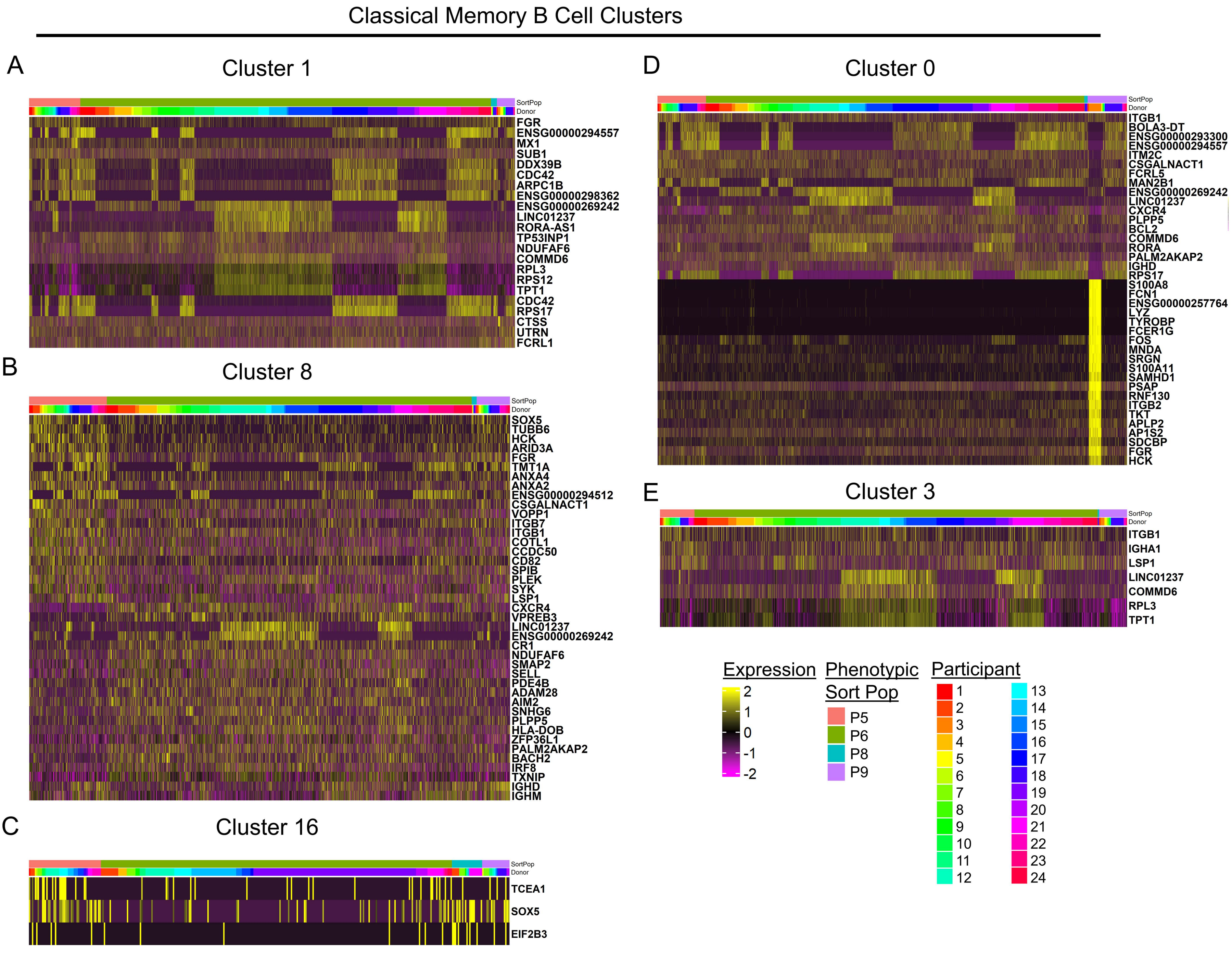

Classical Memory B Cell Clusters
A
Cluster 1
SortPop
Donor
FGR
ENSG00000294557
MX1
SUB1
DDX39B
CDC42
ARPC1B
ENSG00000298362
ENSG00000269242
LINC01237
RORA-AS1
TP53INP1
NDUFAF6
COMMD6
RPL3
RPS12
TPT1
CDC42
RPS17
CTSS
UTRN
FCRL1
B
Cluster 8
SortPop
Donor
SOX5
TUBB6
HCK
ARID3A
FGR
TMT1A
ANXA4
ANXA2
ENSG00000294512
CSGALNACT1
VOPP1
ITGB7
ITGB1
COTL1
CCDC50
CD82
SPIB
PLEK
SYK
LSP1
CXCR4
VPREB3
LINC01237
ENSG00000269242
CR1
NDUFAF6
SMAP2
SELL
PDE4B
ADAM28
AIM2
SNHG6
PLPP5
HLA-DOB
ZFP36L1
PALM2AKAP2
BACH2
IRF8
TXNIP
IGHD
IGHM
C
Cluster 16
SortPop
Donor
TCEA1
SOX5
EIF2B3
D
Cluster 0
SortPop
Donor
ITGB1
BOLA3-DT
ENSG00000293300
ENSG00000294557
ITM2C
CSGALNACT1
FCRL5
MAN2B1
ENSG00000269242
LINC01237
CXCR4
PLPP5
BCL2
COMMD6
RORA
PALM2AKAP2
IGHD
RPS17
S100A8
FCN1
ENSG00000257764
LYZ
TYROBP
FCER1G
FOS
MNDA
SRGN
S100A11
SAMHD1
PSAP
RNF130
ITGB2
TKT
APLP2
AP1S2
SDCBP
FGR
HCK
E
Cluster 3
SortPop
Donor
ITGB1
IGHA1
LSP1
LINC01237
COMMD6
RPL3
TPT1
Expression
2
1
0
-1
-2
Phenotypic Sort Pop
P5
P6
P8
P9
Participant

**Supplemental Figure S8. Differentially expressed genes were identified across phenotypically defined B cell populations within each classical memory B cell cluster.** B cell multi-omics profiling was performed as in **Figure 1** and Methods. The top 20 upregulated genes ($padj < 0.05$, $FC > 1.5$, expressed by >30% of the cells in at least one of the comparator groups, where comparator groups required $n \geq 10$ cells) were identified using the Seurat FindAllMarkers function, with DEGs determined amongst phenotypic populations within each of five transcriptionally defined classical memory B cell clusters. Individual participant ID is also indicated on heatmaps for **A)** Cluster 1, **B)** Cluster 8, **C)** Cluster 16, **D)** Cluster 0, and **E)** Cluster 3.

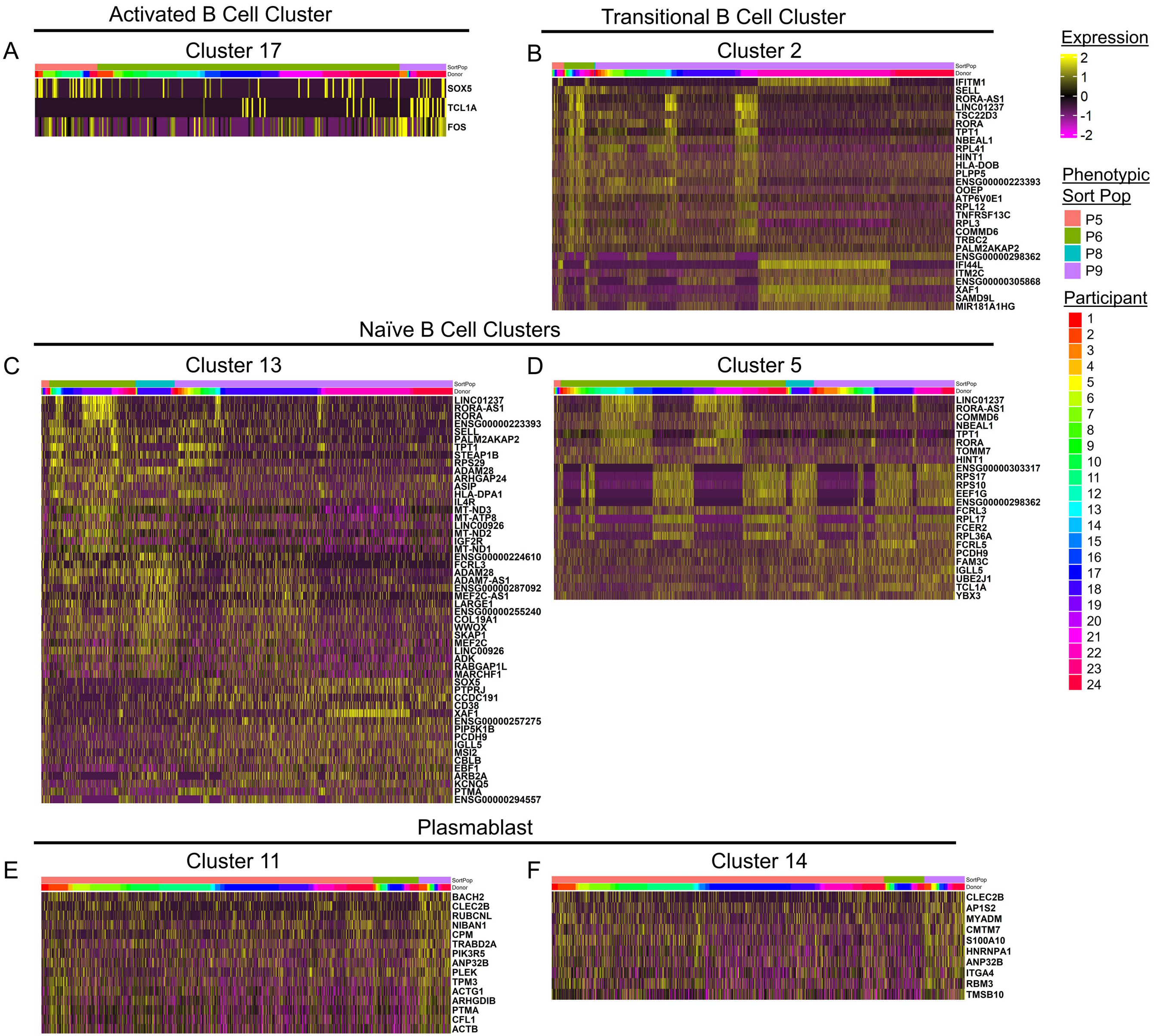

Activated B Cell Cluster
A
Cluster 17
SortPop
Donor
SOX5
TCL1A
FOS
Transitional B Cell Cluster
B
Cluster 2
IFITM1
SELL
RORA-AS1
LINC01237
TSC22D3
RORA
TPT1
NBEAL1
RPL41
HINT1
HLA-DOB
PLPP5
ENSG00000223393
OOEP
ATP6V0E1
RPL12
TNFRSF13C
RPL3
COMMD6
TRBC2
PALM2AKAP2
ENSG00000298362
IFI44L
ITM2C
ENSG00000305868
XAF1
SAMD9L
MIR181A1HG
Expression
2
1
0
-1
-2
Phenotypic Sort Pop
P5
P6
P8
P9
Participant
Naïve B Cell Clusters
C
Cluster 13
LINC01237
RORA-AS1
RORA
ENSG00000223393
SELL
PALM2AKAP2
TPT1
STEAP1B
RPS29
ADAM28
ARHGAP24
ASIP
HLA-DPA1
IL4R
MT-ND3
MT-ATP8
LINC00926
MT-ND2
IGF2R
MT-ND1
ENSG00000224610
FCRL3
ADAM28
ADAM7-AS1
ENSG00000287092
MEF2C-AS1
LARGE1
ENSG00000255240
COL19A1
WWOX
SKAP1
MEF2C
LINC00926
ADK
RABGAP1L
MARCHF1
SOX5
PTPRJ
CCDC191
CD38
XAF1
ENSG00000257275
PIP5K1B
PCDH9
IGLL5
MSI2
CBLB
EBF1
ARB2A
KCNQ5
PTMA
ENSG00000294557
D
Cluster 5
LINC01237
RORA-AS1
COMMD6
NBEAL1
TPT1
RORA
TOMM7
HINT1
ENSG00000303317
RPS17
RPS10
EEF1G
ENSG00000298362
FCRL3
RPL17
FCER2
RPL36A
FCRL5
PCDH9
FAM3C
IGLL5
UBE2J1
TCL1A
YBX3
Plasmablast
E
Cluster 11
BACH2
CLEC2B
RUBCNL
NIBAN1
CPM
TRABD2A
PIK3R5
ANP32B
PLEK
TPM3
ACTG1
ARHGDIB
PTMA
CFL1
ACTB
F
Cluster 14
CLEC2B
AP1S2
MYADM
CMTM7
S100A10
HNRNPA1
ANP32B
ITGA4
RBM3
TMSB10

**Supplemental Figure S9. Differentially expressed genes were identified across phenotypically defined B cell populations within individual activated, transitional, naïve, and plasmablast clusters.** B cell multi-omics profiling was performed as in **Figure 1** and Methods. The top 20 upregulated genes (padj < 0.05, FC > 1.5, expressed by >30% of the cells in at least one of the comparator groups, where comparator groups required n ≥ 10 cells) were identified using the Seurat FindAllMarkers function, with DEGs determined amongst phenotypic populations within each transcriptionally defined B cell cluster. Individual participant ID is also indicated on heatmaps for **A)** Activated Cluster 17, **B)** Transitional Cluster 2, **C)** Naïve Cluster 13, **D)** Naïve Cluster 5, **E)** Plasmablast Cluster 11, and F) Plasmablast Cluster 14.

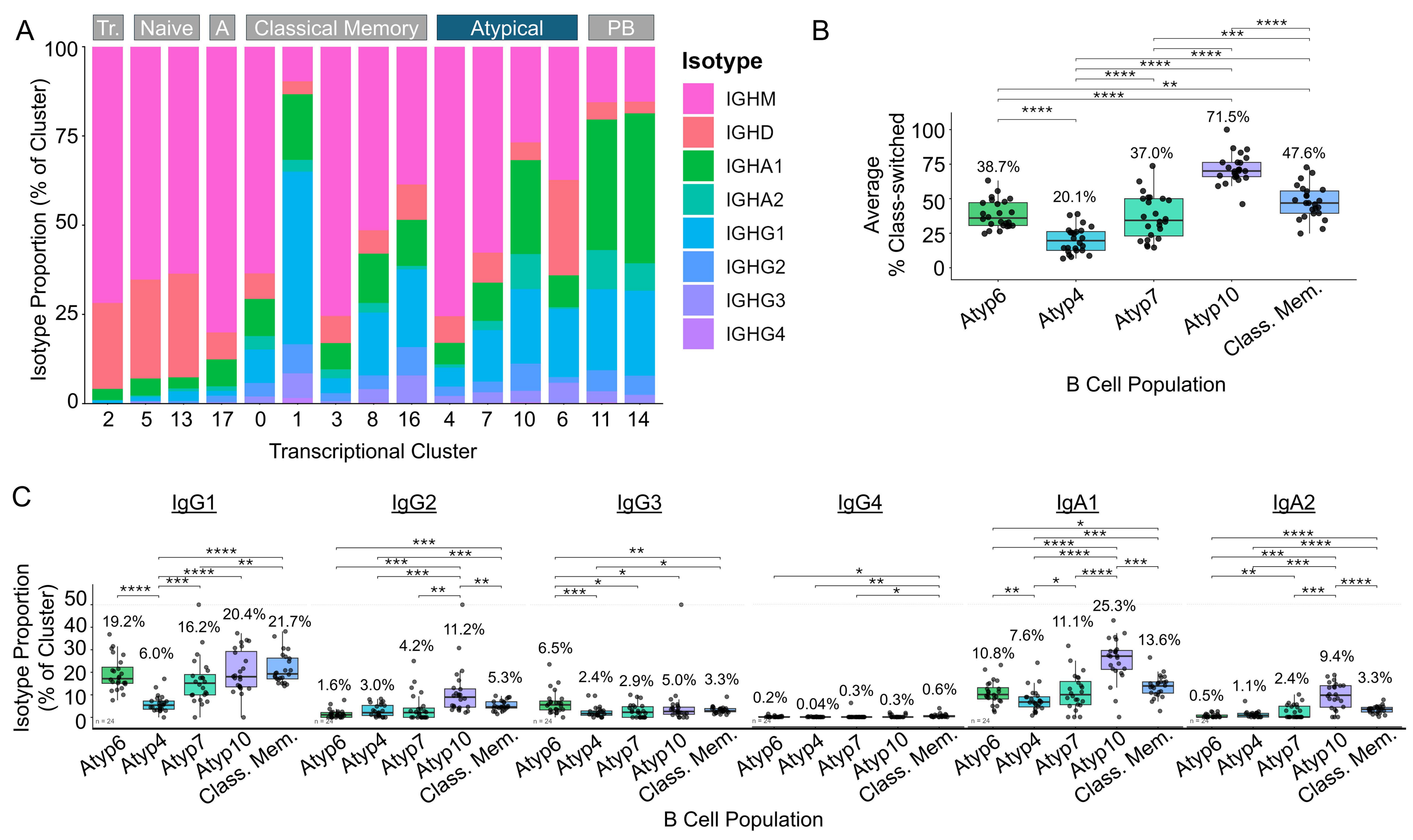

A
Tr.
Naive
A
Classical Memory
Atypical
PB
Isotype Proportion (% of Cluster)
Transcriptional Cluster
Isotype
IGHM
IGHD
IGHA1
IGHA2
IGHG1
IGHG2
IGHG3
IGHG4
B
Average % Class-switched
38.7%
20.1%
37.0%
71.5%
47.6%
Atyp6
Atyp4
Atyp7
Atyp10
Class. Mem.
B Cell Population
C
IgG1
IgG2
IgG3
IgG4
IgA1
IgA2
Isotype Proportion (% of Cluster)
19.2%
6.0%
16.2%
20.4%
21.7%
1.6%
3.0%
4.2%
11.2%
5.3%
6.5%
2.4%
2.9%
5.0%
3.3%
0.2%
0.04%
0.3%
0.3%
0.6%
10.8%
7.6%
11.1%
25.3%
13.6%
0.5%
1.1%
2.4%
9.4%
3.3%
n = 24
B Cell Population

**Supplemental Figure S10. Differences in class switching are observed between atypical B cell populations.** B cell multi-omics profiling was performed as in **Figure 1** and Methods. **A)** The proportion of each isotype within each cluster is shown. **B-C)** Individual participants are plotted (n = 24). **B)** The frequency of isotype-switched (IgA+ or IgG+) B cells is shown within each of the indicated populations. **C)** Frequencies of each switched isotype were calculated within each of the indicated populations. In panel B-C, Freidman test with paired Wilcoxon signed-rank test (Benjamin Hochberg correction), **** $p < 0.0001$, *** $p < 0.001$, ** $p < 0.01$, * $p < 0.05$, all other comparisons are not significant.

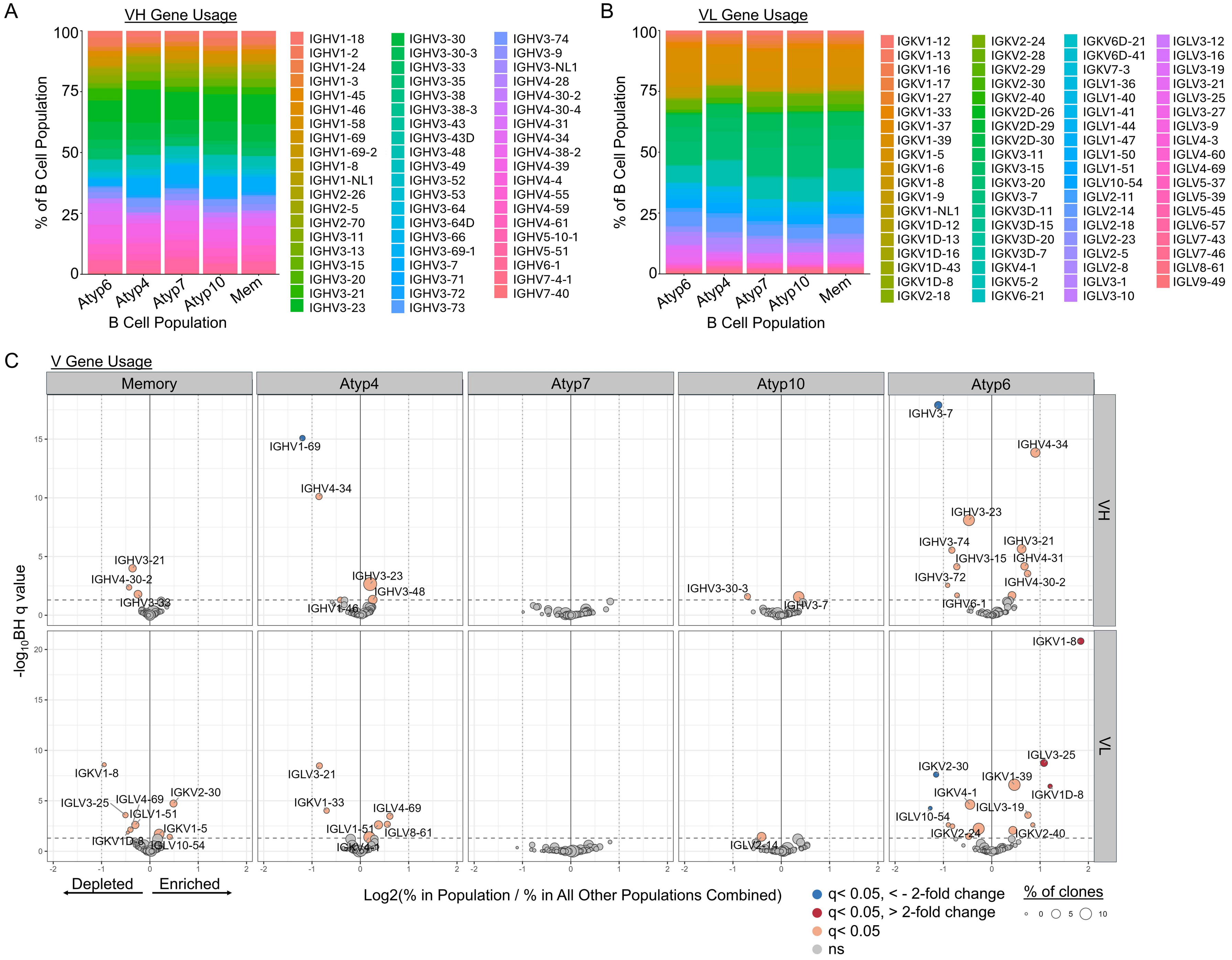

A
VH Gene Usage
% of B Cell Population
B Cell Population
Atyp6
Atyp4
Atyp7
Atyp10
Mem
IGHV1-18
IGHV1-2
IGHV1-24
IGHV1-3
IGHV1-45
IGHV1-46
IGHV1-58
IGHV1-69
IGHV1-69-2
IGHV1-8
IGHV1-NL1
IGHV2-26
IGHV2-5
IGHV2-70
IGHV3-11
IGHV3-13
IGHV3-15
IGHV3-20
IGHV3-21
IGHV3-23
IGHV3-30
IGHV3-30-3
IGHV3-33
IGHV3-35
IGHV3-38
IGHV3-38-3
IGHV3-43
IGHV3-43D
IGHV3-48
IGHV3-49
IGHV3-52
IGHV3-53
IGHV3-64
IGHV3-64D
IGHV3-66
IGHV3-69-1
IGHV3-7
IGHV3-71
IGHV3-72
IGHV3-73
IGHV3-74
IGHV3-9
IGHV3-NL1
IGHV4-28
IGHV4-30-2
IGHV4-30-4
IGHV4-31
IGHV4-34
IGHV4-38-2
IGHV4-39
IGHV4-4
IGHV4-55
IGHV4-59
IGHV4-61
IGHV5-10-1
IGHV5-51
IGHV6-1
IGHV7-4-1
IGHV7-40
B
VL Gene Usage
% of B Cell Population
B Cell Population
IGKV1-12
IGKV1-13
IGKV1-16
IGKV1-17
IGKV1-27
IGKV1-33
IGKV1-37
IGKV1-39
IGKV1-5
IGKV1-6
IGKV1-8
IGKV1-9
IGKV1-NL1
IGKV1D-12
IGKV1D-13
IGKV1D-16
IGKV1D-43
IGKV1D-8
IGKV2-18
IGKV2-24
IGKV2-28
IGKV2-29
IGKV2-30
IGKV2-40
IGKV2D-26
IGKV2D-29
IGKV2D-30
IGKV3-11
IGKV3-15
IGKV3-20
IGKV3-7
IGKV3D-11
IGKV3D-15
IGKV3D-20
IGKV3D-7
IGKV4-1
IGKV5-2
IGKV6-21
IGKV6D-21
IGKV6D-41
IGKV7-3
IGLV1-36
IGLV1-40
IGLV1-41
IGLV1-44
IGLV1-47
IGLV1-50
IGLV1-51
IGLV10-54
IGLV2-11
IGLV2-14
IGLV2-18
IGLV2-23
IGLV2-5
IGLV2-8
IGLV3-1
IGLV3-10
IGLV3-12
IGLV3-16
IGLV3-19
IGLV3-21
IGLV3-25
IGLV3-27
IGLV3-9
IGLV4-3
IGLV4-60
IGLV4-69
IGLV5-37
IGLV5-39
IGLV5-45
IGLV6-57
IGLV7-43
IGLV7-46
IGLV8-61
IGLV9-49
C
V Gene Usage
Memory
Atyp4
Atyp7
Atyp10
Atyp6
VH
VL
-log10BH q value
Log2(% in Population / % in All Other Populations Combined)
Depleted
Enriched
q< 0.05, < - 2-fold change
q< 0.05, > 2-fold change
q< 0.05
ns
% of clones

**Supplemental Figure S11. Distinct atypical B cell populations show differences in VH and VL gene usage.** B cell transcriptional and immune repertoire profiling was performed as in **Figure 1** and Methods. **A)** VH and **B)** VL gene usage is shown within atypical and classical memory B cell populations **C)** VH (*top*) and VL (*bottom*) enrichment or depletion at the clonotype level was calculated for each atypical or classical memory B cell population compared to the remaining atypical/memory populations pooled, as in Methods. Enrichment/depletion was measured as log2 fold change of % V gene in each B cell population / % V gene in remaining populations and calculated by Fisher Exact test with Benjamin Hochberg correction, q value < 0.05. Color of circle corresponds to significance and fold change cutoffs. Size of circle corresponds to % of clones expressing V gene.

A

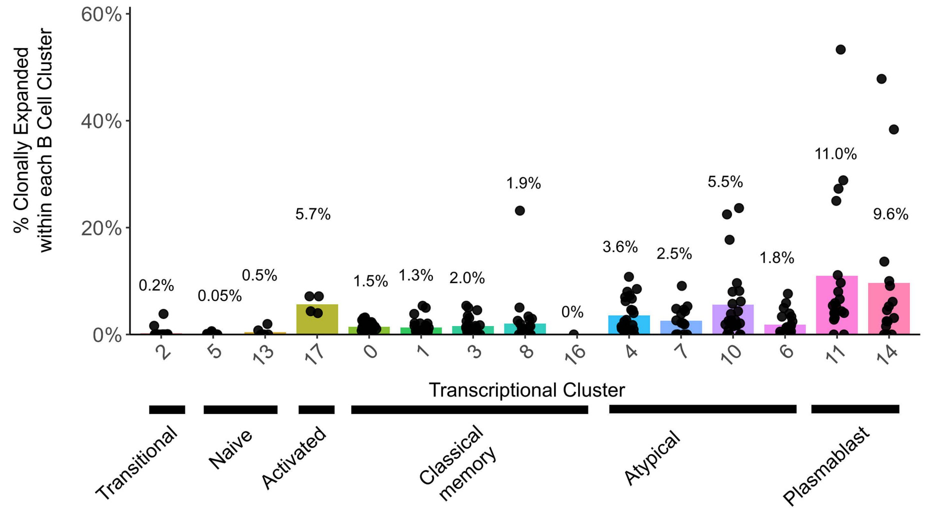

| Cluster | Participants (n) | Cells (n) |
|---|---|---|
| 2 | 23 | 7994 |
| 5 | 23 | 5824 |
| 13 | 6 | 365 |
| 17 | 4 | 129 |
| 0 | 24 | 10026 |
| 1 | 24 | 8014 |
| 3 | 24 | 6396 |
| 8 | 23 | 2472 |
| 16 | 1 | 25 |
| 4 | 23 | 6080 |
| 7 | 16 | 838 |
| 10 | 23 | 1954 |
| 6 | 22 | 3654 |
| 11 | 19 | 1907 |
| 14 | 15 | 1290 |

B

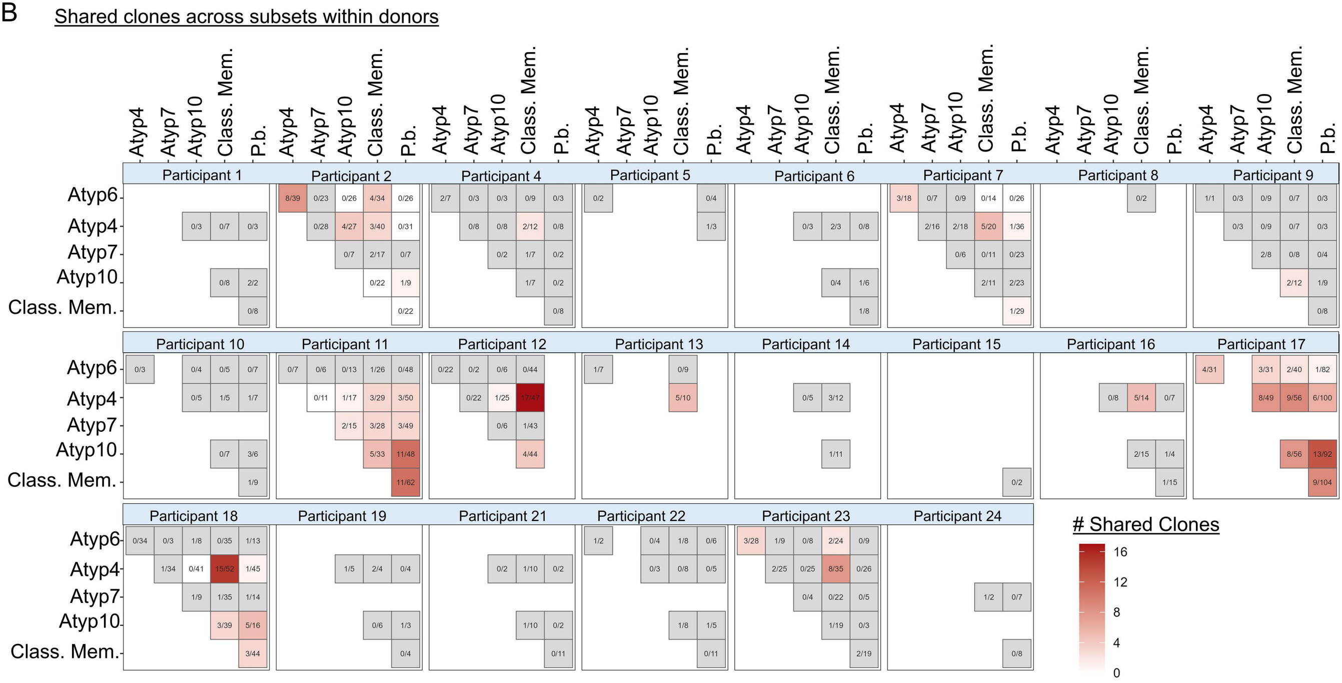

**Supplemental Figure S12. Clonal expansion varies across transcriptionally defined clusters, and across individuals, with clonal overlap observed between atypical, classical memory, and plasmablast B cells in most individuals.** B cell transcriptional and immune repertoire profiling was performed as in **Figure 1** and Methods. **A)** The frequency of clonally expanded cells (at least two clones per clonotype within an individual) was calculated and is plotted for individual participants within B cell clusters, with means indicated above each bar (left). Table (right) indicates number of participants and total cells per cluster. Cell populations with n < 20 cells were excluded. **B)** Clonal overlap is depicted, with the scale representing the number of clones shared between the indicated B cell populations within each participant. The number of shared clones is listed within each box, with comparisons of n < 5 clones in grey boxes. Cell populations with n < 20 cells, singlets, and any individuals that did not have at least two subsets that met these criteria were excluded from this analysis.
